# Cik1 and Vik1 Accessory Proteins Confer Distinct Functions to the Kinesin-14, Kar3

**DOI:** 10.1101/2022.09.09.507361

**Authors:** Zane J Bergman, Jonathan J Wong, David G Drubin, Georjana Barnes

## Abstract

The budding yeast *Saccharomyces cerevisiae* has a closed mitosis in which the mitotic spindle and cytoplasmic microtubules (MTs) used to segregate chromosomes remain separated by the nuclear envelope throughout the cell cycle. Kar3, the yeast kinesin-14, has unique roles in both compartments and has been implicated in capturing unattached kinetochores, stabilizing crosslinked interpolar microtubules (MT), and creating intranuclear and cytoplasmic MT arrays at the spindle pole body for kinetochore capture and karyogamy, respectively. Here, we show that two proteins, Cik1 and Vik1, that form heterodimers with Kar3, regulate its localization and function within the cell and along MTs in a cell cycle-dependent manner. Using a cell cycle synchronized, yeast MT dynamics reconstitution assay in cell lysate, we found that Kar3Vik1 induces MT catastrophes in S phase and metaphase and limits MT polymerization in G1 and anaphase. In contrast, Kar3Cik1 is a catastrophe and pause promoter in G1, while increasing catastrophes in metaphase and anaphase. Adapting this assay to track single-molecules, we saw that Kar3Cik1 is necessary for tracking MT plus-ends in S phase and metaphase, but, surprisingly, not during anaphase. These experiments demonstrate how the binding partners of Kar3 modulate its diverse functions both spatially and temporally.

**SUMMARY STATEMENT:** We show through biochemical reconstitution experiments and live-cell imaging that the functions and localization of the budding yeast kinesin-14, Kar3, are dictated by which of its two accessory protein binding partners, Cik1 or Vik1, it binds to and by the cell cycle stage.

## INTRODUCTION

Microtubules are highly dynamic polymers with myriad functions in cells including providing mechanical support for large cellular structures, acting as tracks for intracellular cargo, and forming the mitotic spindle to faithfully segregate chromosomes to daughter cells (Chaaban and Brouhard, 2017). These different MT forms and functions are achieved through a host of microtubule-associated proteins (MAPs) that control MT interactions and assembly dynamics. MAPs can affect MT assembly and disassembly through various mechanisms including as polymerases, depolymerases, stabilizers, and as destabilizers. One particularly important class of MAPs is the motor proteins, which generate forces on MTs for moving cargos within the cell, separating spindle poles and chromosomes, and for bending cilia and flagella. The concerted effects of these many different types of MAPs ultimately dictate MT form and function. In the budding yeast, *Saccharomyces cerevisiae*, MTs have one known essential function in vegetatively growing cells: to faithfully segregate sister chromatids to daughter cells during mitosis. The various steps of this complex process, which include spindle assembly and movement to the bud neck, and chromosome segregation, occur concurrently in budding yeast due, in large part, to the distinct properties imparted by MAPs independently in the nucleus and cytoplasm. These functions are kept separate throughout the cell cycle due to a closed mitosis. The MAPs responsible for this regulation are either completely excluded from one compartment, or are post-translationally modified differentially across compartments, to give rise to these different functions in the nucleus and in the cytoplasm.

In the case of Kar3, previous studies established that its subcellular localization is determined by its binding partner, of which there are two. The Kar3Cik1 heterodimer is transported into the nucleus because of a nuclear localization sequence in Cik1 (Manning et al., 1999). In the nucleus, Kar3Cik1 generates spindle forces (Hoyt et al., 1993; Saunders et al., 1997a), properly arrays interpolar MTs (Molodtsov et al., 2016), has a role in kinetochore capture (Liu et al., 2011; Middleton and Carbon, 1994), and transports newly-captured chromosomes along MTs (Tanaka et al., 2005; Tanaka et al., 2007). The Kar3Vik1 heterodimer is only cytoplasmic (Manning et al., 1999). The functions of this complex are presently not known. Previous work showed that Kar3 absence increases the number and length of cytoplasmic MTs (Huyett et al., 1998; Saunders et al., 1997b), suggesting a role in assembly regulation, and provided evidence that Kar3Vik1 focuses MT minus ends on the cytoplasmic face of the spindle pole body (SPB), the budding yeast MT organizing center (King et al., 2021).

Dissecting the functions specific to each type of Kar3 heterodimer provides a unique opportunity to investigate a novel mode of kinesin regulation, but has been challenging. The activity of Kar3 heterodimers on MTs was first described as being minus-end-directed translocation with low processivity (Endow et al., 1994), a property common to other studied kinesin-14 family members. Subsequent Kar3 studies provided evidence for its potential to switch between plus- and minus-end-directed movement (Molodtsov et al., 2016), or to stably associate with a growing plus end (Sproul et al., 2005), possibly when in complex with Bim1, the budding yeast EB1 homolog (Kornakov et al., 2020; Ogren et al., 2022; Sproul et al., 2005). In addition, while early work with Kar3 provided evidence for plus-end depolymerase activity (Sproul et al., 2005), later work failed to document this activity (Mieck et al., 2015).

The *in vitro* assays of Kar3 function and regulation cited above provided insights into possible Kar 3 functions, but the inconsistency of the results also is a source of confusion. While these studies were pioneering efforts to elucidate Kar3 function, they might have suffered from complicating issues such as use of heterogeneous assay constituents and innovative but nevertheless non-physiological recombinant fusion proteins to stably form each of the two Kar3 heterodimers (Chu et al., 2005; Rank et al., 2012). Those earlier reconstitution assays might also have suffered from lack of the context of post-translational modifications present within native yeast cells. Importantly, these modifications are likely to be specific to the cell cycle phase. Also missing from previous *in vitro* MT dynamics studies using purified proteins is the complete panoply of MAPs present within the cell, and thus the possibility to uncover emergent properties of the system in its full complexity.

We previously developed a biochemical reconstitution assay that harnesses the genetic tractability of budding yeast and enables imaging of individual MTs, which is very difficult in living cells (Bergman et al., 2019). In this assay, dynamics of single MTs are observed using TIRF microscopy. Moreover, the assay uses total soluble cellular protein, and is performed in a cell-cycle specific context by preparing lysates from cells arrested at specific cell cycle stages. Here, we utilize this *in vitro* assay to reveal the biochemical activities of the two distinct Kar3 heterodimers on MT dynamics in the context of the complexity of the cell at each stage of the cell cycle. We also describe the association of Kar3 heterodimers with, and locomotion along, MTs. In total, we show that Kar3Cik1 and Kar3Vik1 have similar but distinct motility profiles and functions that are regulated in the cell cycle and are intrinsic to the respective heterodimer, and not simply a product of their subcellular localization. Complementary *in vivo* studies test the applicability of these results to live cells.

## RESULTS

### Kar3 Heterodimer Localization Is Cell-cycle Dependent

Because subcellular localization provides important information about biological function, we set out to more fully describe the localization of the two types of Kar3 heterodimers through the cell cycle. We generated strains that express a GFP-Kar3 fusion protein from the native *KAR3* locus. These strains also express *mRUBY2-TUB1*, for visualizing MTs, and temperature-sensitive *cdc* alleles, that allow arrest at different cell cycle stages. The strains were arrested at either G1 (*cdc28-4*), S phase (*cdc7-1*), metaphase (*cdc23-1*), or late anaphase (*cdc15-2*) (Goranov et al., 2009). Arrested cells were observed by time-lapse, wide-field fluorescence microscopy at the restrictive temperature (Fig. 1A). Cells arrested in G1 had the stereotypical bright nuclear microtubule bundle and long cytoplasmic MTs (cMTs) with a single GFP-Kar3 spot at the spindle pole body (SPB). When cells were arrested in S phase, the majority of cells had short spindles and long cMTs. GFP-Kar3 had a bright spot at the SPBs, and appeared along the spindle, and as a lobed haze that is typical of kinetochore localization (Xiangwei et al., 2000). For cells arrested in metaphase, short spindles were fully covered in GFP-Kar3. We also observed, at very low frequency, GFP spots on the plus ends of cMTs. Presumably, these spots represent Kar3Vik1 molecules, which previously had only been described at the cytoplasmic face of the SPB. The SPB’s also showed bright GFP-Kar3 signals. During anaphase, GFP-Kar3 was observed at the SPBs, along the length of the spindle, and as a lobed haze close to the SPB, again likely reflecting kinetochore localization.

**Figure 1.**
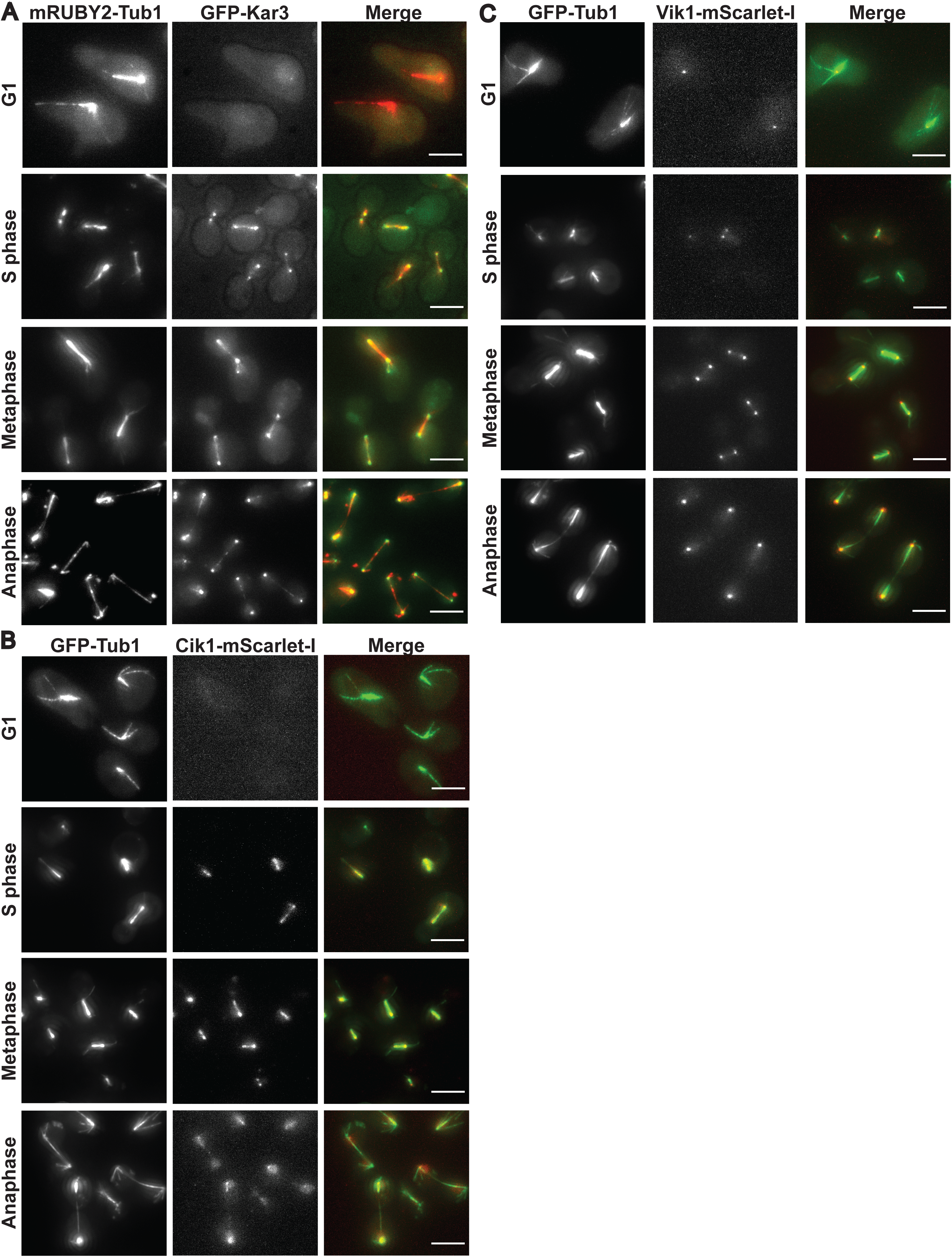
Cell-cycle-dependent Localization of Kar3 Heterodimers. **(A)** GFP-Kar3 localization along mRUBY2-Tub1 MTs at either G1, S phase, metaphase, or anaphase. **(B)** Cik1-mScarlet-I localization along GFP-Tub1 MTs throughout the cell cycle. **(C)** Vik1-mScarlet-I localization along GFP-Tub1 MTs throughout the cell cycle. Scale bars are 5 µm.

We then determined the localization of the distinct Kar3 heterodimers by tagging either Cik1 or Vik1 with mScarlet-I paired with *GFP-TUB1*, and arresting the cells using conditional *cdc* alleles, as above. Of immediate interest, we found that Cik1 was not detectable in G1-arrested cells (Fig. 1B). Conversely, Vik1 localized to SPBs (Fig. 1C), indicating that the single spot seen at the SPB in G1-arrested GFP-Kar3 cells consisted of only Kar3Vik1 heterodimers. At metaphase, Cik1 was present on short spindles and at SPBs while Vik1 was observed on the spindle poles at SPBs, presumably on the cytoplasmic SPB face. Upon arrest in metaphase, Cik1 localized in a bi-lobed configuration along short spindles. This pattern is typical of proteins associated with kinetochores. Vik1-mScarlet-I was found mainly on the cytoplasmic side of SPBs, but with a slight haze of signal between the SPBs. Noticeably absent was any detectable Vik1 signal on the plus-ends of cMTs like we observed in the GFP-Kar3 cells. Cik1 localization in anaphase-arrested cells was observed along the extended spindles and as a haze near the SPBs in the interior of the nucleus, typical of kinetochores. Some cells contained bright puncta at SPBs. Vik1 localization during anaphase was very similar to what was observed during metaphase. During this stage of the cell cycle, there was a bright spot on the cytoplasmic face of the SPB surrounded by a faint haze. These observations reaffirm that Kar3 localization changes in a programmed manner throughout the vegetative cell cycle and is dependent upon its binding partner, with Cik1 transporting the kinesin into the nucleus via its nuclear-localization sequence and Vik1 retaining it in the cytoplasm.

Next, we wanted to investigate the roles of Cik1 or Vik1 in MT organization and assembly in cells and for Kar3 localization throughout the cell cycle. To achieve this, we introduced either a *CIK1-AID* or *VIK1-AID* allele into the *GFP-KAR3 mRUBY2-TUB1 cdc^ts^* strains used above. The *AID* motif is an auxin-inducible degron that triggers proteolysis of the tagged protein in the presence of 3-indole acetic acid (IAA). The AID-tagged proteins were depleted for 30 min from cells that had been at restrictive temperature for 3 hours. Western blot analysis showed that this was sufficient time for these proteins to be depleted by over 90%. These strains gave us tight control over when proteins were absent and therefore helped prevent unwanted secondary effects of the absence of these proteins. When cells were mounted on coverslips, the pre-warmed media also contained IAA to continue the depletion during the observation time. Cells arrested in G1 and depleted of Cik1 retained a single spot of GFP-Kar3 at SPBs as was seen in non-depleted cells (Fig. 2A). When cells arrested in S phase were depleted of Cik1, GFP-Kar3 was not observed in the nucleus. During metaphase in cells depleted of Cik1, GFP-Kar3 was absent from the nucleus, but a strong GFP-Kar3 signal was observed on the cytoplasmic face of the SPB. The increase in Kar3 on cMTs could be due to more Vik1 binding to Kar3 in the absence of Cik1. Additionally, GFP-Kar3 was absent from the short kinetochore microtubules (kMTs).

**Figure 2.**
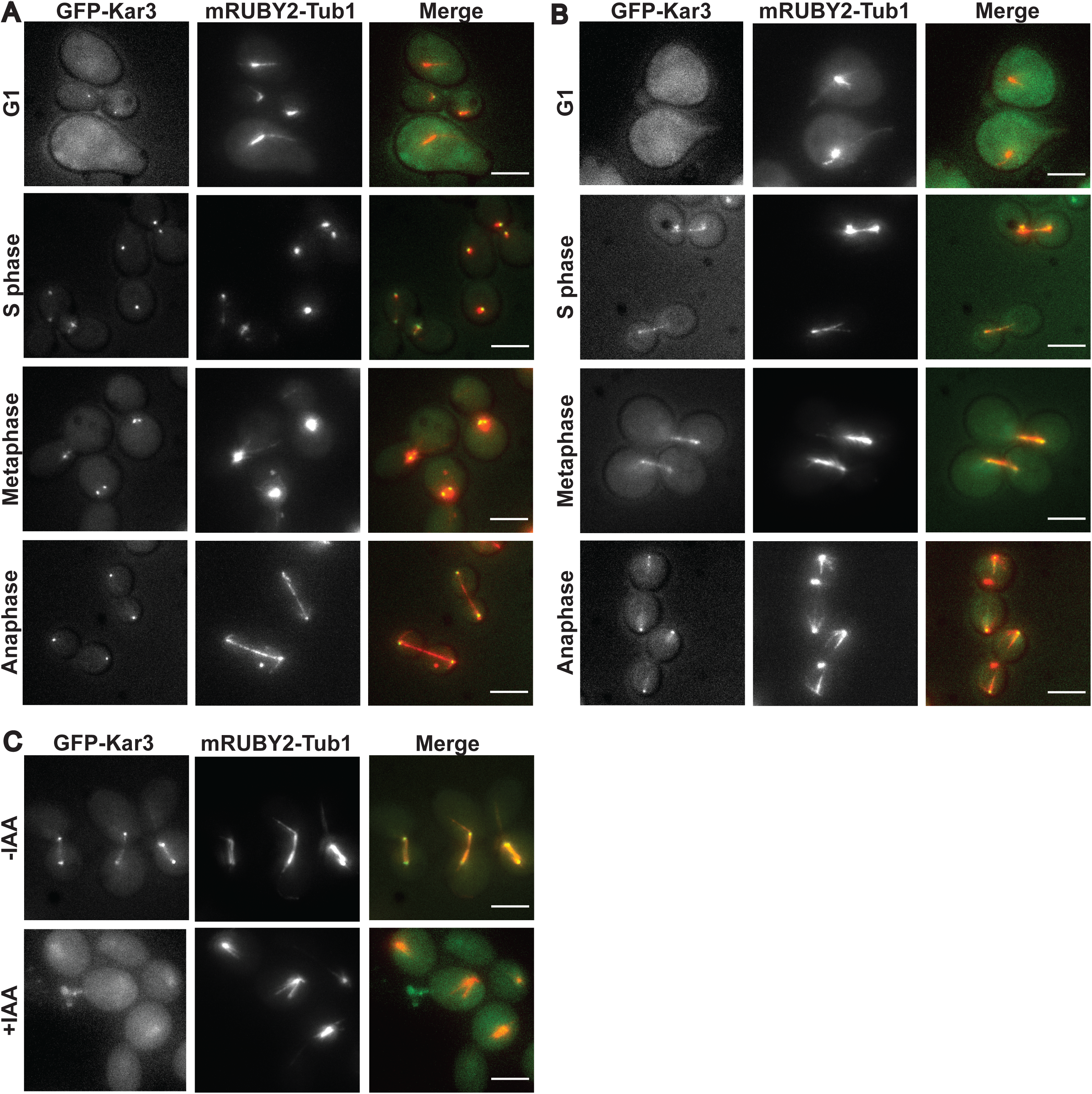
Kar3 Accessory Proteins Direct Kar3 Localization and MT Morphology. **(A)** GFP-Kar3 localization and mRUBY2-Tub1 morphology in cell-cycle-arrested cells depleted of Cik1-AID. **(B)** GFP-Kar3 localization and mRUBY2-Tub1 morphology in cell-cycle-arrested cells depleted of Vik1-AID. **(C)** GFP-Kar3 and mRUBY2-Tub1 localization in cells with CIK1-AID VIK1-AID alleles. Top row has not been treated with IAA. Bottom row has been treated with IAA for 30 min. Scale bars are 5 µm.

Depleting Vik1 from cells confirmed the difference in heterodimer localization. During G1, Vik1-depleted cells lacked the GFP-Kar3 spot at the SPB seen in WT and Cik1-AID cells (Fig. 2B). GFP-Kar3 had a reduced signal at the SPBs in S phase-arrested cells depleted of Vik1. During metaphase and upon depletion of Vik1, GFP-Kar3 was detected inside the nucleus at the SPB and along the spindle. In anaphase-arrested cells depleted of Vik1, GFP-Kar3 was absent from the cytoplasm but present on the nuclear face of SPBs with a cloud of signal near them, consistent with kinetochore localization.

We were curious to determine if Kar3 is able to interact with MTs in cells in the absence of both binding partners, Cik1 and Vik1. We created a strain that had both *CIK1-AID* and *VIK1-AID* alleles along with *GFP-KAR3* and *mRUBY2-TUB1*. This strain also contained the *cdc23-1* allele to arrest cells in metaphase. When these cells were arrested and the Kar3 binding partners depleted, as above, no GFP-Kar3 could be detected at the SPBs or in association with the nucleus (Fig 2C). To test the converse possibility, that Cik1 or Vik1 relied on Kar3 for their interaction with MTs, we introduced a *KAR3-AID* allele into our *GFP-TUB1* strains that had either *CIK1* or *VIK1* tagged with mScarlet-I. Upon depletion of Kar3 from these strains, the mScarlet signal became diffuse throughout the cell and there were no puncta, even at the SPBs (Fig. S1). These data demonstrate that only heterodimers, and not individual Kar3, Cik1 or Vik1 proteins, can associate with MTs *in vivo*.

### Kar3Cik1 Controls Spindle Length While Kar3Vik1 Affects Cytoplasmic MTs

Previous studies described MT morphology phenotypes for *kar3Δ* cells that typically include longer, more numerous cMTs (Huyett et al., 1998; Saunders et al., 1997b) and short or broken metaphase spindles (Hoyt et al., 1993; Saunders et al., 1997a). Similar phenotypes were observed in single-depletion cells, which we here set out to define quantitatively according to cell cycle stage. Most notable were differences in spindle lengths between the cells. For measuring spindles, GFP-TUB1 cells were arrested in either metaphase or anaphase and were then depleted of either Kar3, Cik1, or Vik1 using the AID system. Cells were then observed under the arrest and depletion conditions. Measurement of spindles was done at the first time point. In Kar3-AID and Cik1-AID cells arrested in metaphase, spindles could form, but some spindles would break. During time-lapse observation, 29.8% of intact spindles in Kar3-depleted and 36.5% in Cik1-depleted cells would break. In Kar3- and Cik1-depleted cells the mean distance between SPBs of intact spindles was much shorter compared to non-depleted cells (1.6 ± 0.9, 1.5 ± 0.8, and 2.8 ± 1.1μm, respectively) (Fig. 3A). Spindles in Vik1-depleted metaphase-arrested cells were similar in length to WT spindles (2.2 ± 0.6 μm) and did not exhibit breaking spindles. To further characterize this phenotype, again, we utilized a strain that could simultaneously deplete Cik1 and Vik1. The spindles in these metaphase cells were on average 1.3 ± 0.6 μm long and resembled those of Kar3-AID and Cik1-AID cells. The above observations are consistent with those in previous reports on spindles in *kar3*Δ, *cik1*Δ, *and vik1*Δ cells (Hepperla et al., 2014), which indicate that the Kar3Cik1 motor is needed to both balance forces along the interpolar MTs and to capture kinetochores during metaphase. As for cells arrested in late anaphase and depleted of Kar3, their spindle lengths were similar to WT (7.7 ± 1.1 and 7.8 ± 1.2 μm, respectively) (Fig. 3B), possibly indicating that Kar3 heterodimers are not necessary for maintaining elongated spindles. However, the picture becomes more complex when either Cik1 or Vik1 is depleted. When Cik1 was depleted, elongated spindles were longer than WT (10.1 ± 3.2 μm) and occasionally exceeded the length of the cell, bending around the cell interior. Cells arrested in late anaphase and depleted of Vik1 also had elongated spindles (8.9 ± 1.9 μm average). Since these two accessory proteins are in different cellular compartments and the magnitude of this increase is different, the mechanism of increasing the length of spindles most likely differs.

**Figure 3.**
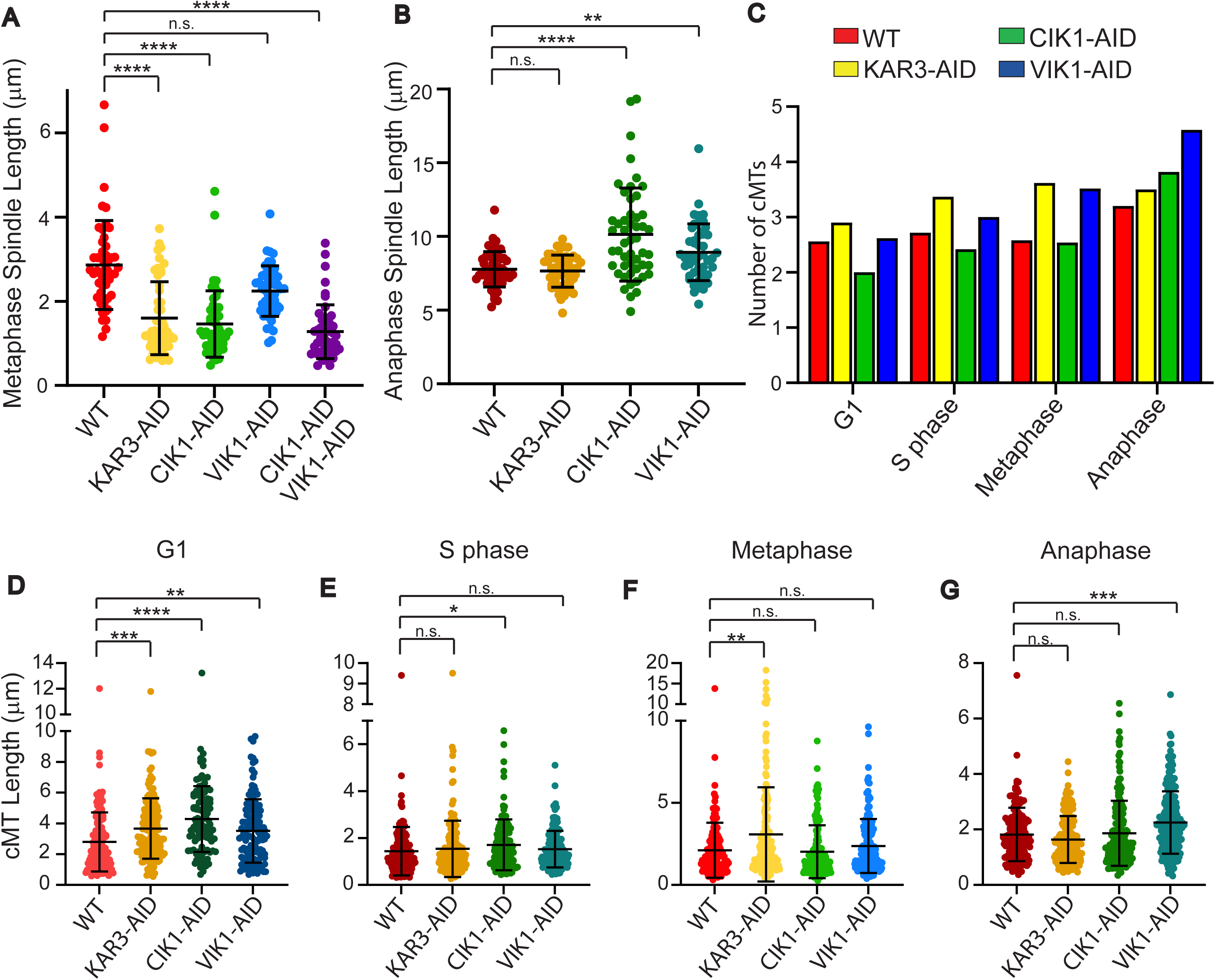
Spindle Length and cMT Morphology in Depleted Cells. **(A)** Spindle length of cells arrested in metaphase and either otherwise WT or depleted of Kar3, Cik1, Vik1, or Cik1 and Vik1. n= 50. **(B)** Spindle lengths in cells arrested in late anaphase that are either WT or depleted of Kar3, Cik1, or Vik1. n= 50. **(C)** Average number of cMTs per cell during the cell cycle in either WT (red), Kar3-(yellow), Cik1-(green), or Vik1-depleted (blue) cells. n= 50. The lengths of all cMTs from 50 cells that are either WT, Kar3-, Cik1, or Vik1-depleted and arrested at either G1 **(D)**, S phase **(E)**, metaphase **(F)**, or anaphase **(G)**. Designations for statistics are as follows: ns = not significant, * = p<0.05, ** = p<0.01, *** = p<0.0005, **** = p<0.0001.

Previous research found that cells lacking functional Kar3 have more cMTs and that these cMTs are longer on average than those observed in WT cells (Huyett et al., 1998; Saunders et al., 1997b). We quantified for different cell cycle stages both cMT number and length in strains that expressed *GFP-TUB1*, had an AID-tagged protein, and that carried conditional-lethal *CDC* mutant alleles so they could be arrested at different cell cycle stages. We found that cells depleted of either Kar3 or Vik1 had an increased number of cMTs at all cell cycle stages (Fig. 3C) as compared to WT. Only during late anaphase did Cik1-depleted cells have more cMTs than WT. Interestingly, the lengths of cMTs in Kar3- and Vik1-depleted cells were almost indistinguishable from WT during all cell cycle stages except G1, where all depletions increased the number of cMTs (Figs.3D-F). Interestingly, Vik1-depletion was the only condition to have increased cMT length in late anaphase. (Fig. 3G).

### Kar3 Heterodimers’ Distinct Contributions to MT Dynamics Vary with the Cell Cycle Stage

Knowing that Kar3Cik1 and Kar3Vik1 have distinct subcellular localizations, we set out to compare their biochemical activities. Previous work with Kar3 revealed a potential MT depolymerase activity and possible interactions with MAPs that are known to regulate MT dynamics (Endow et al., 1994; Sproul et al., 2005; Tirnauer et al., 1999). We therefore set out to determine the effects of the two different Kar3 heterodimers on MT dynamics in our previously developed reconstitution assay (Bergman et al., 2019). This assay enables observation of dynamics of single MTs in the presence of total soluble protein lysate.

Earlier work from our lab using this reconstitution assay showed that the effects of Kar3Cik1 and Kar3Vik1 heterodimers on MT dynamics are quite different. Lysates created from asynchronous WT cultures do not assemble MTs. A lack of Cik1 in lysate from asynchronous cells did not change this result. In contrast, lysates lacking Vik1 could readily polymerize MTs (Bergman et al., 2019). We investigated the contributions of each heterodimer to MT dynamics regulation further and with greater precision by utilizing our *GFP-TUB1*-expressing strains that had either *KAR3*, *CIK1*, or *VIK1* tagged with an AID motif on their C-terminus. These strains also each contained a temperature-sensitive allele of a *CDC* gene to enable cells to be arrested at different cell cycle stages. We then prepared whole cell lysates from these arrested and depleted cultures and assayed MT dynamics in them.

Unlike lysates from otherwise WT cells arrested in G1 with the *cdc28-4* temperature-sensitive allele in which MTs do not assemble, lysates depleted of Kar3, Cik1, or Vik1 can each assemble dynamic MTs. MTs in lysates made from G1-arrested Kar3-depleted cells assembled MTs at a rate of 0.38 ± 0.16 µm min^-1^ (Fig. 4A), with a disassembly rate of 0.78 ± 0.37 µm min^-1^ (Fig. 4B). MTs assembled faster (0.66 ± 0.17 and 0.52 ± 0.20 µm min^-1^, respectively) in Cik1-depleted and Vik1-depleted lysates.

**Figure 4.**
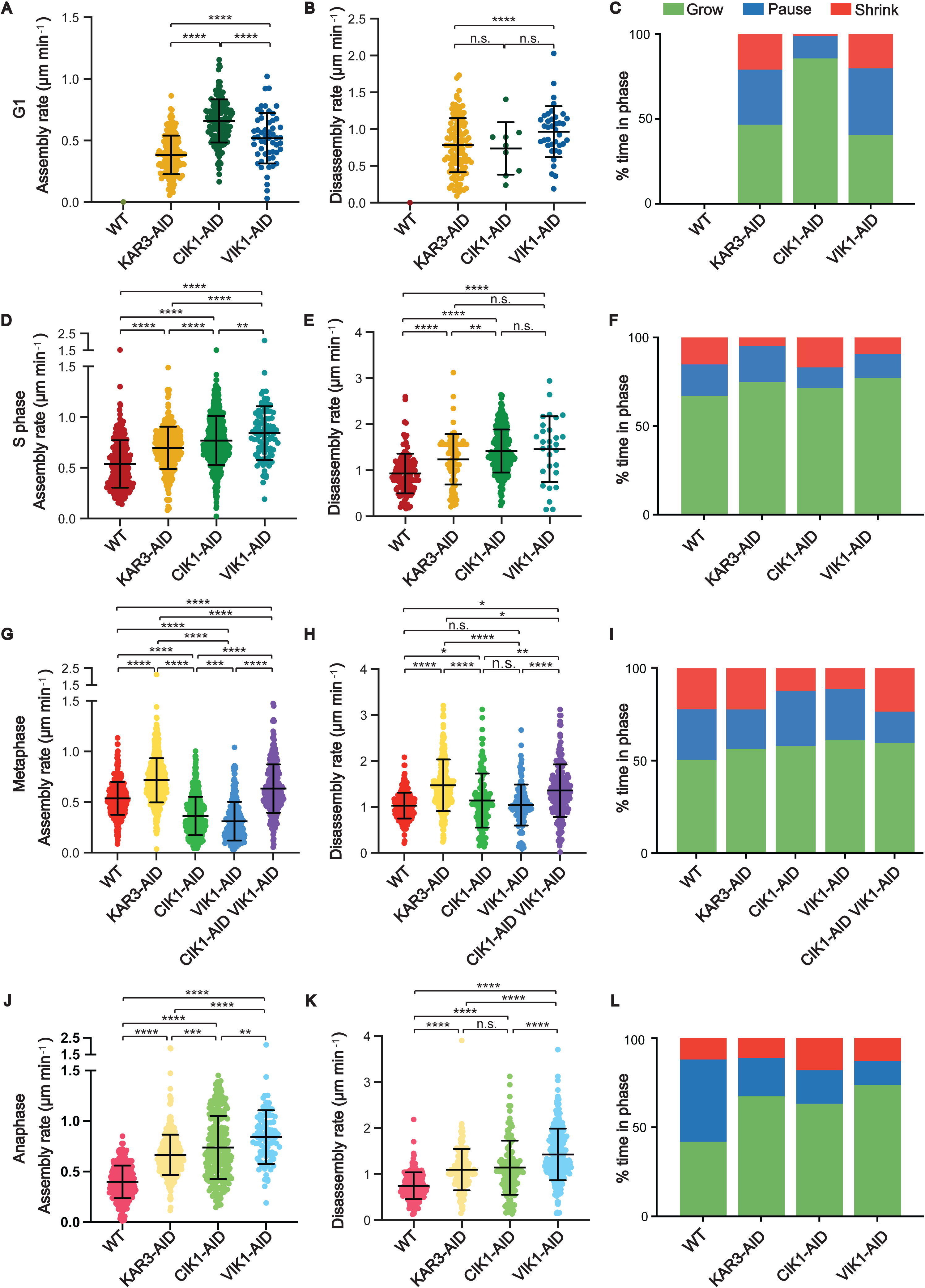
Kar3, Cik1, and Vik1 Have Different Effects on Reconstituted MT Dynamics. Reconstituted MT dynamics in lysates from cells arrested in either G1, S phase, metaphase, or anaphase were quantified. Differences in MT assembly rates **(A, D, G, and J)**, disassembly rates **(B, E, H, and K)** or growth parameters **(C, F, I, and L)** were seen between WT, Kar3-, Cik1-, and Vik1-depleted lysates. Designations for statistics are as follows: ns = not significant, * = p<0.05, ** = p<0.01, *** = p<0.0005, **** = p<0.0001.

These MTs also disassembled faster (0.74 ± 0.36 and 0.97 ± 0.35 µm min^-1^, respectively). Globally, MTs in Kar3-depleted lysates grew 46.6% of the time, shrank 21% of the time, and paused for the remaining 32.4% of the time (Fig. 4C). MTs in Cik1-depleted lysates grew 85.6% of the time and only paused 1.2% of the time. Surprisingly, MTs in Vik1-depleted lysates only grew 40.7% of the time and shrank 20.2% of the time, resembling the distribution seen in Kar3-depleted lysates. The frequency of catastrophe in Kar3- and Vik1-depleted lysates was about 10-fold higher than that of MTs in the Cik1-depleted lysate (2.7 or 3.1 vs 0.3 x10^-3^ events sec^-1^) (Table 1). These data suggest that in G1, Kar3 heterodimers either behave as MT depolymerases or they prevent MT polymerization via an unknown mechanism. This conclusion is in agreement with previously reported data (Huyett et al., 1998; Saunders et al., 1997b; Tirnauer et al., 1999) and our own cellular imaging (Fig. 3D), which show an increase in the number and length of cMTs in unbudded cells lacking Kar3. These data further suggest that Kar3Cik1 heterodimers may be the main depolymerase or catastrophe factor that prevents MTs from growing in G1-arrested lysates.

**Table 1.** Parameters of MT Dynamics in G1-arrested Lysates.

| G1-arrested | WT | KAR3-AID<br>n = 35 | CIK1-AID<br>n = 34 | VIK1-AID<br>n = 12 |
| --- | --- | --- | --- | --- |
| Assembly rate<br>($\mu\text{m min}^{-1}$) | - | $0.38 \pm 0.16$ (169) | $0.66 \pm 0.17$ (146) | $0.52 \pm 0.20$ (54) |
| Disassembly rate<br>($\mu\text{m min}^{-1}$) | - | $0.78 \pm 0.37$ (122) | $0.74 \pm 0.36$ (9) | $0.97 \pm 0.34$ (38) |
| Freq. of Cat.<br>($\text{sec}^{-1}$)$\times 10^3$ | - | 2.7 | 0.3 | 3.1 |
| Freq. of Rescue<br>($\text{sec}^{-1}$)$\times 10^3$ | - | 9.5 | 23.4 | 11.9 |
| Dynamicity<br>(dimers $\text{sec}^{-1}$) | - | 9.0 | 16.0 | 11.0 |

In S phase-arrested lysates depleted of Kar3, Cik1, or Vik1, MT assembly and disassembly rates both increased relative to rates in lysates from S phase-arrested wild-type cells (Fig. 4D, E). The overall proportion of time when MTs were growing increased for all three lysates relative to the proportion in lysates from WT cells (Fig. 4F). Thus, in S phase-arrested lysates, Kar3 heterodimers appear to be responsible for muting assembly and disassembly rates. Interestingly, the frequency of catastrophe was significantly less than in WT lysates in only the Kar3- and Vik1-depleted lysates (Table 2). This result was reflected in the amount of time MTs in these lysates spent shrinking (Fig. 4F). Thus, Kar3Vik1 acts to promote catastrophes during S phase.

**Table 2.** Parameters of MT Dynamics in S phase-arrested Lysates.

| S phase<br>-arrested | WT<br>n = 47 | KAR3-AID<br>n = 65 | CIK1-AID<br>n = 123 | VIK1-AID<br>n = 18 |
| --- | --- | --- | --- | --- |
| Assembly rate<br>($\mu\text{m min}^{-1}$) | $0.54 \pm 0.23$ (248) | $0.70 \pm 0.21$ (280) | $0.77 \pm 0.24$ (612) | $0.84 \pm 0.26$ (275) |
| **Disassembly rate**<br>($\mu\text{m min}^{-1}$) | $0.93 \pm 0.43$ (131) | $1.24 \pm 0.55$ (77) | $1.42 \pm 0.47$ (318) | $1.46 \pm 0.71$ (29) |
| **Freq. of Cat.**<br>($\text{sec}^{-1}$)$\times 10^3$ | 2.0 | 0.8 | 1.9 | 1.1 |
| **Freq. of Rescue**<br>($\text{sec}^{-1}$)$\times 10^3$ | 11.0 | 14.9 | 8.6 | 8.9 |
| **Dynamicity**<br>(dimers $\text{sec}^{-1}$) | 13.1 | 16.6 | 22.8 | 22.8 |

Upon acute Kar3, Cik1, or Vik1 depletion in lysates from metaphase-arrested cells via the AID degron system, it became clear that Kar3Cik1 and Kar3Vik1 perform different functions in MT dynamics regulation. When Kar3 is depleted from metaphase lysates, MT assembly (0.54 ± 016 vs 0.72 ± 0.22 µm min^-1^) and disassembly (1.03 ± 0.28 vs 1.51 ± 0.91 µm min^-1^) rates both increase relative to wild-type (Fig. 4G). Interestingly, the assembly rates in the Cik1- and Vik1-depleted metaphase lysates were slower than in WT lysates (Fig. 4G), while disassembly rates were similar across all 3 conditions (Fig. 4H). MTs in the Kar3-depleted lysates spend less time paused than MTs in WT lysates (27% WT; 21% Kar3-AID) (Fig. 4I), while MTs in either Cik1- or Vik1-depleted lysates have about 70% of the frequency of catastrophe compared to MTs in WT lysates (Table 3). This lowered catastrophe rate leads to a decrease in time MTs spend shrinking compared to MTs in WT lysates (Figure 3I). We further investigated this phenomenon using the strain with both *CIK1* and *VIK1* tagged with AID. In lysates created from this strain and arrested in metaphase, MT dynamics were similar to those in Kar3-depleted lysates in that both assembly and disassembly rates were faster than in WT lysates and in that the frequency of catastrophe and dynamics were increased (Fig. 4G, H, and Table 1). These data suggest that, during metaphase, Kar3Cik1 and Kar3Vik1 regulate MT dynamics by separate mechanisms.

**Table 3.**
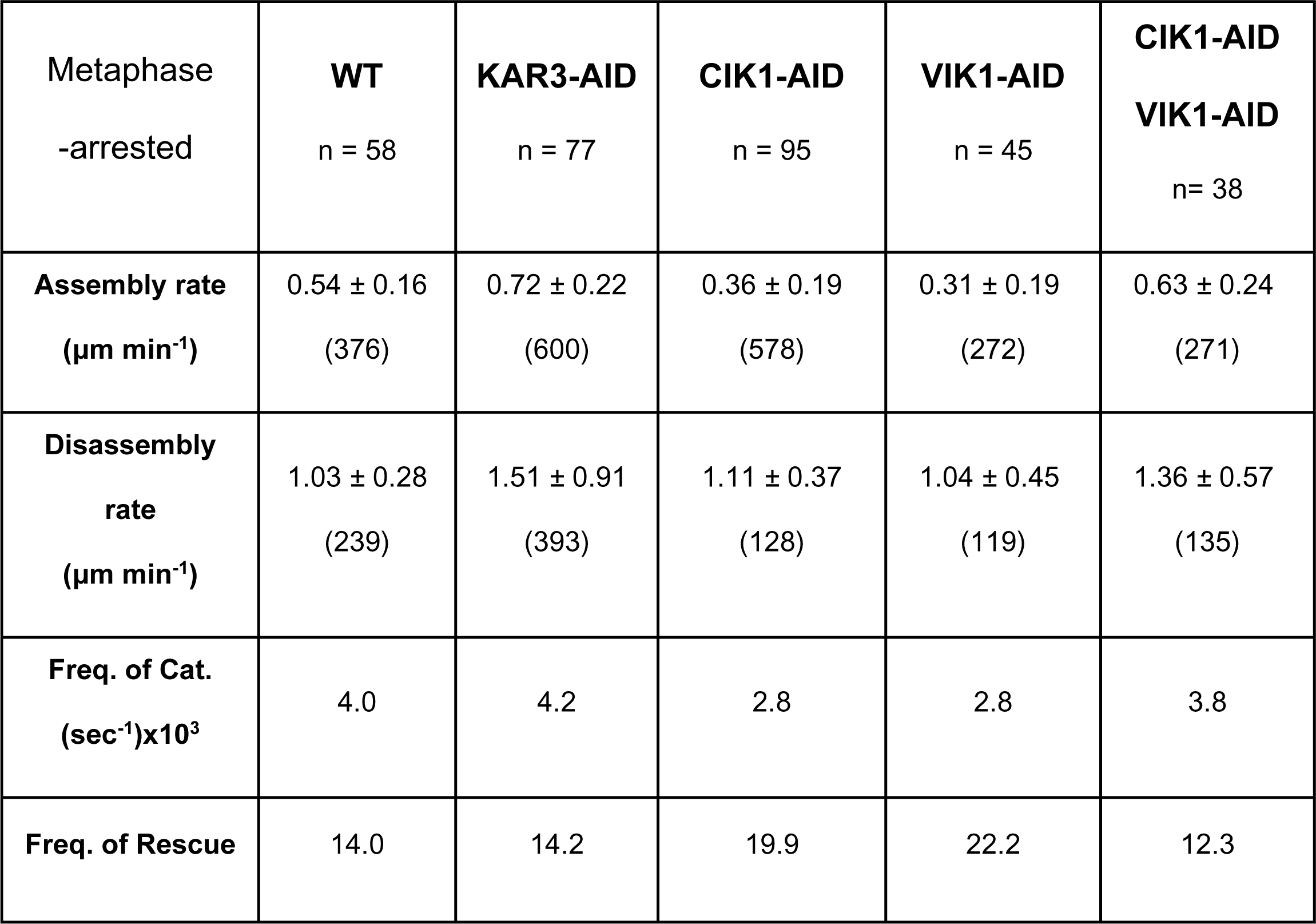
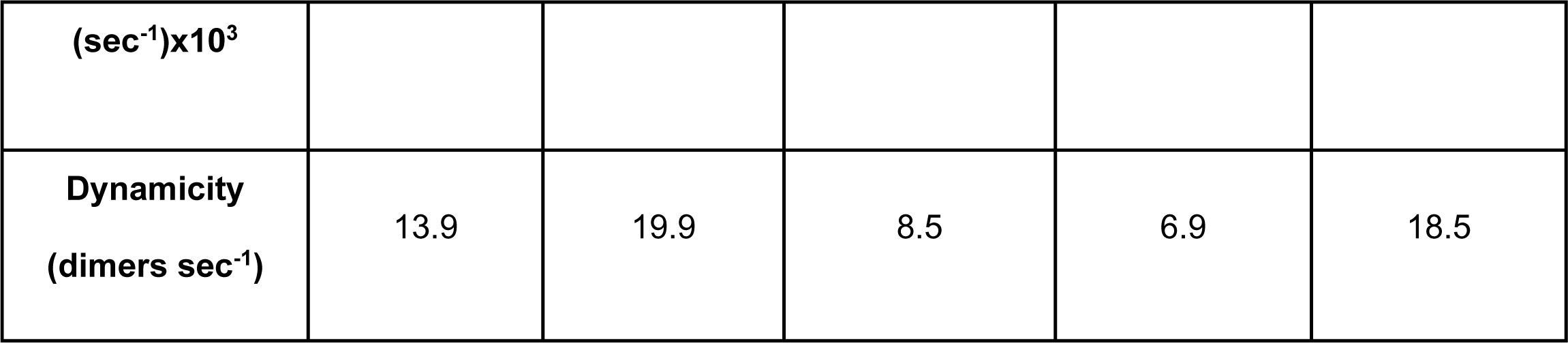
Parameters of MT Dynamics in metaphase-arrested Lysates.

The apparent functions of Kar3Cik1 and Kar3Vik1 heterodimers diverge most sharply in anaphase. When MT dynamics parameters are measured in WT, *KAR3-AID*, *CIK1-AID*, and *VIK1-AID* lysates made from anaphase-arrested cells, a complex picture emerges. All depletions increase both the MT assembly and disassembly rates (Fig. 4J, K). And all of the depletions decrease the time MTs spend paused, as compared to MTs in WT lysates (Fig. 4L), making for highly dynamic MTs. However, in the Kar3- and Cik1-depleted lysates, MTs had similar disassembly rates and catastrophe frequency. However, the MTs in Vik1-depleted lysate are even faster and more dynamic (Table 4). These data suggest that Kar3 heterodimers normally act to increase MT pausing in anaphase and that Kar3Vik1 has a stronger effect than Kar3Cik1. These results mirror MT behavior observed in late anaphase cells, wherein cMTs and kMTs are short and stable.

**Table 4.** Parameters of MT Dynamics in anaphase-arrested Lysates.

| Anaphase<br>-arrested | **WT**<br>n = 105 | **KAR3-AID**<br>n = 82 | **CIK1-AID**<br>n = 52 | **VIK1-AID**<br>n = 113 |
| --- | --- | --- | --- | --- |
| **Assembly rate**<br>**(µm min<sup>-1</sup>)** | 0.51 ± 0.25 (391) | 0.67 ± 0.20 (394) | 0.74 ± 0.31 (244) | 0.80 ± 0.28 (91) |
| **Disassembly rate**<br>**(µm min<sup>-1</sup>)** | 0.93 ± 0.48 (193) | 1.09 ± 0.45 (157) | 1.14 ± 0.59 (128) | 1.43 ± 0.56 (239) |
| **Freq. of Cat.**<br>**(sec<sup>-1</sup>)x10<sup>3</sup>** | 2.3 | 1.4 | 1.5 | 1.5 |
| **Freq. of Rescue**<br>**(sec<sup>-1</sup>)x10<sup>3</sup>** | 16.3 | 10.9 | 6.1 | 9.1 |
| **Dynamicity**<br>**(dimers sec<sup>-1</sup>)** | 10.8 | 15.5 | 19.0 | 22.6 |

### Kar3 Heterodimers Have Disparate Associations with MTs

To determine if each of the two Kar3 heterodimers has a unique localization or dynamic profile on MTs independent of intracellular localization, we created soluble, whole-cell lysates from strains expressing *GFP-KAR3* and *mRUBY2-TUB1* that could be arrested at different cell cycle stages using a temperature-sensitive allele of a *CDC* gene. From lysates created from metaphase-arrested cells, we observed four different classes of GFP-Kar3 interactions with MTs polymerized from HiLyte647-labeled tubulin seeds: tracking the plus-end of a growing MT (Fig. 5A), interaction with the stable minus-end of a seed (Fig. 5B), movement along the MT lattice toward the plus-end (Fig. 5C), and movement along the lattice toward the MT minus-end (Fig. 5D).

**Figure 5.**
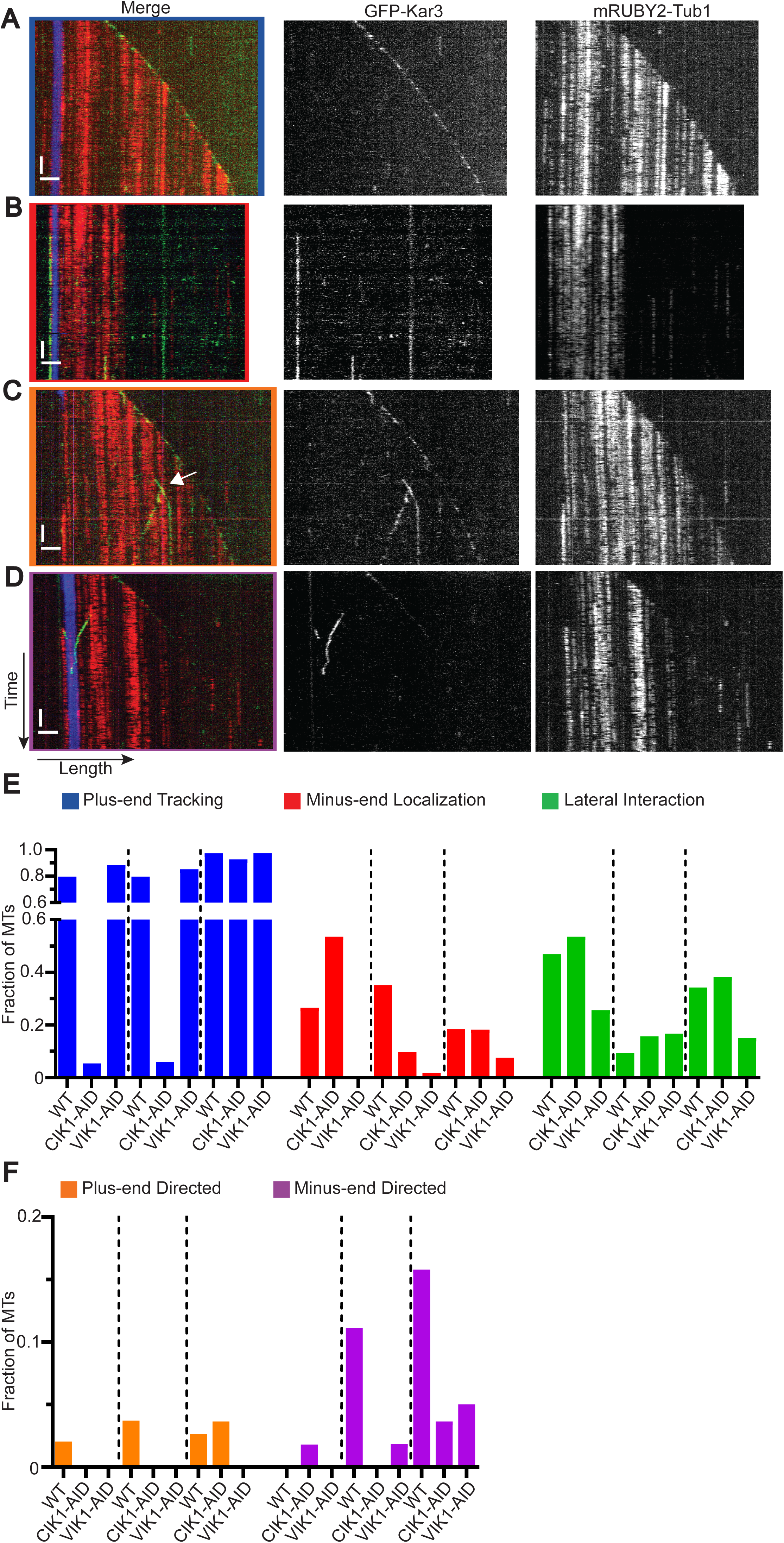
GFP-Kar3 Heterodimers’ Different Associations with Reconstituted MTs. Dynamic MTs were reconstituted from lysate from cells expressing GFP-KAR3 (green) and mRUBY2-TUB1 (red). These MTs were adhered to coverslips using HiLyte647-labeled seeds (blue). Kymographs were made to observe motor association and translocation along the MT. **(A)** GFP-Kar3 tracking the growing plus-end of a MT. **(B)** GFP-Kar3 interacting with the minus end of a MT seed. **(C)** GFP-Kar3 processively moving toward the plus-end of a growing MT (white arrow). **(D)** GFP-Kar3 processively moving toward the minus-end of a MT. White horizontal scale bars are 2 µm and white vertical bars are 2 min. Fraction of observed MTs that display non-motile **(E)** and motile **(F)** interactions with GFP-Kar3. Dashed lines separate cell cycle stages S phase, Metaphase, Anaphase. Color outlines of merged images in A-D correspond to bars in E and F.

We next set out to determine which heterodimer showed each behavior during different cell cycle phases. We analyzed soluble lysates that lacked either Cik1 or Vik1 by use of the AID system and counted the number of times we saw these different interactions (Table 5). During S phase, in WT lysates, 79.6% of MTs had a GFP-Kar3 on the growing plus-end of MTs. This number dropped to only 5.4% in Cik1-depleted lysates and increased to 88.8% of MTs in Vik1-depleted lysates, implicating Cik1 in plus-end tracking during S phase. Interestingly, the amount of GFP-Kar3 found on the minus-ends of seeds increased in Cik1-depleted lysates (53.6%) compared to WT (33.3%) and this localization was absent in Vik1-depleted lysates. This observation matches our *in vivo* data (Fig. 2B) wherein GFP-Kar3 does not have strong SPB signal in S phase-arrested, Vik1-depleted cells. Also of note was that static interaction of GFP-Kar3 along the MT lattice dropped from around half of observed MTs in WT and Cik1-depleted lysates to only 25.6% of MTs in Vik1-depleted lysates.

**Table 5.**
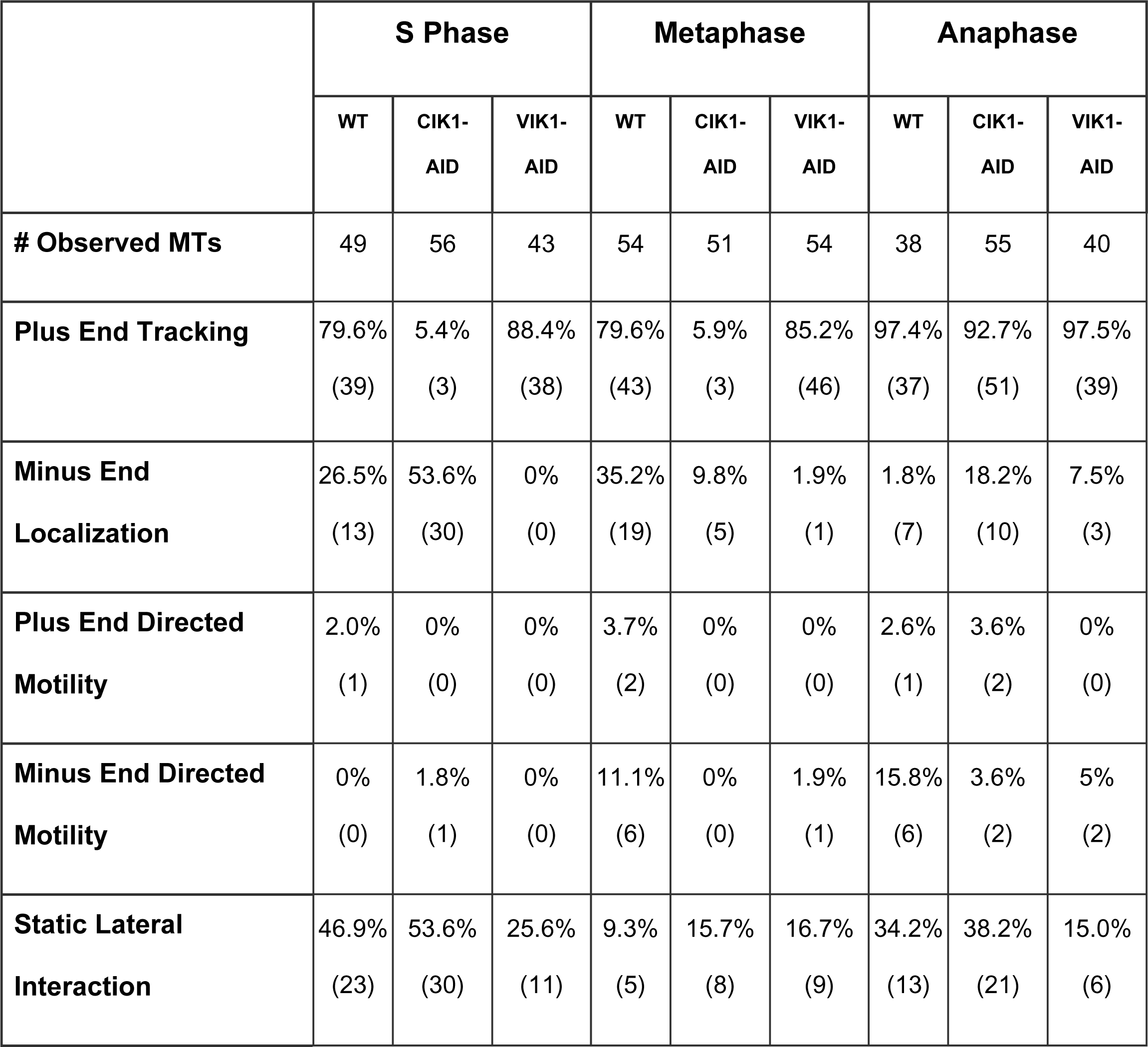
GFP-Kar3 Motility Observed on Single MTs.

In metaphase-arrested lysates, GFP-Kar3 was observed tracking the plus-end at some point in 79.6% of observed MTs. This number dropped to only 5.9% of MTs when Cik1 was depleted. However, GFP-Kar3 was observed tracking plus-ends in 85.2% of MTs in Vik1-depleted lysates. These observations demonstrate that Cik1 is necessary for Kar3 to track plus-ends during metaphase as well. GFP-Kar3 interacted with the stabilized MT minus-end in 35.2% of MTs in WT extracts. The instances of these interactions decreased to 9.8% of MTs in Cik1-depleted lysates and 1.9% in Vik1-depleted lysates. This observation matches with our *in vivo* data showing that GFP-Kar3 localizes with both sides of the SPB, from which yeast MTs. GFP-Kar3 no longer localized to the SPB on the nuclear or cytoplasmic side of the nuclear envelope when the corresponding Kar3 accessory protein was depleted. We observed minus-end directed motility of Kar3 on 11.1% of MTs in lysates from WT cells. Motility was not observed in the Cik1-depleted lysates. Only 1.9% of the MTs in Vik1-depleted lysates had motors moving in the minus-end direction. These data suggest that the majority of minus-end-directed motility we observed in our assay is dependent upon Kar3Cik1. Interestingly, in both Cik1 and Vik1 depletions, the number of MTs with static, lateral interactions with GFP-Kar3 increased as compared to WT.

Once again, in anaphase, the activities of the two heterodimers appear to diverge from each other. Surprisingly, Cik1 depletion did not prevent GFP-Kar3 from tracking the plus-end, or from translocating toward the plus end along the lattice. In contrast, Vik1 depletion decreased the number of MTs with GFP-Kar3 on the minus-end, minus-end-directed motility, and static interactions along the lattice. Additionally, GFP-Kar3 was now observed diffusely along lengths of MTs in Cik1-depleted lysates during anaphase although the function of this is uncertain (Fig. S2). These and our other results point to an important role for Vik1 during anaphase.

We also took this opportunity to measure the rate of Kar3 minus-end directed motility. We determined the rate of movement in a manner similar to calculating the rates of MT assembly and disassembly. In our assay, GFP-Kar3 moves toward the minus end at a rate of 0.64 ± 0.27 μm min^-1^ (10.6 nm min^-1^) in metaphase-arrested lysate, which is much slower than previously reported rates (Allingham et al., 2007; Chu et al., 2005; Endow et al., 1994; Mieck et al., 2015). However, this rate is similar to the rate reported for captured centromeres that move along the lattice of kMTs toward the minus-end in a Kar3-dependent manner (Tanaka et al., 2007). The processivity or GFP-Kar3 in our assay was low, with run lengths that averaged only 0.88 ± 0.57 μm.

## DISCUSSION

In this work we used *in vivo* studies and our previously developed reconstitution system (Bergman et al., 2019) to examine in detail the localization and functions of the budding yeast kinesin-14, Kar3, and its two binding partners, Cik1 and Vik1, and how these change through the cell cycle. During mitotic growth these three proteins form two distinct heterodimers, Kar3Cik1 and Kar3Vik1. The former is found exclusively in the nucleus, and the latter is found exclusively in the cytoplasm (Manning et al., 1999). By expressing these proteins, along with tubulin, as fluorescent protein fusions in cells carrying temperature-sensitive alleles of cell cycle control genes, we were able to describe their localization during four distinct cell cycle stages. Then, by selectively disrupting one heterodimer at a time using an inducible degradation system for either Cik1 or Vik1, we were able to confirm the localization pattern specific to each heterodimer.

### Accessory Factors Cik1 and Vik1 Impart Distinct Functionality to Kar3

We also determined the contribution of these heterodimers to overall MT dynamics regulation using our reconstitution system. We found that though these heterodimers are very similar in structure, they make distinct contributions to MT dynamics regulation at distinct cell cycle stages. We further investigated the mechanisms by which Kar3Cik1 and Kar3Vik1 heterodimers interact with MTs to produce these effects by adapting our reconstitution assay to follow single molecules in three-color TIRF microscopy. We observed complex Kar3 behaviors: motility along MTs, static interactions with either the minus-end or the lattice, and tracking of dynamic plus-ends. Interestingly, these activities were dependent on which binding partner was associated with Kar3 and to the stage of the cell cycle. In total, these studies determined where the two different Kar3 heterodimers are located during each cell cycle stage and how they affect MT dynamics, providing a more complete understanding of how they produce their unique functions.

Our previous reconstitution studies provided evidence for differences in the functions of the two Kar3 heterodimers. While WT lysates from cells arrested in G1 did not assemble MTs, similarly arrested lysates from *vik1Δ* cells could assemble MTs, though lysates from *cik1Δ* cells in G1 could not (Bergman et al., 2019). This observation motivated us to answer outstanding questions about the function and regulation of Kar3 in budding yeast.

Our *in vitro* studies showed that Kar3 heterodimers induce catastrophes while dampening the overall MT assembly and disassembly rates (Fig. 4 and Tables 1-4). When the gene encoding Cik1 or Vik1 is deleted, the catastrophe frequency decreases and the overall dynamicity of MTs increases because assembly and disassembly rates increase. Interestingly, the Kar3Cik1 heterodimer appears to control the number of catastrophes during G1, whereas Kar3Vik1 has a more prominent role in S phase. Both heterodimers contribute to dynamicity and catastrophe frequency during mitotic phases.

Previous studies have indicated that Kar3Cik1 actively catalyzes MT depolymerization (Endow et al., 1994; Sproul et al., 2005), while Kar3Vik1 does not (Allingham et al., 2007). A number of untested mechanisms have been proposed for how Kar3 moves from the MT plus-end and destabilizes the MT as it translocates towards the minus end (Hou and Wang, 2010). Our single-molecule dynamics data suggest a higher level of functional and mechanistic complexity. Though we observed GFP-Kar3 molecules tracking disassembling plus-ends, GFP-Kar3 was also found on growing plus-ends and moving along the lattice toward the minus-end without causing catastrophes (Fig. 5, Table 5). The fact that Kar3 could have disparate depolymerase activity in differently prepared lysates (Fig. 4F, I, and L and Fig. 5E) indicates that more work is needed to fully elucidate both its the mechanism and regulation. The observation that Kar3 depletion increases both assembly and disassembly rates is a strong indication that further studies are warranted. Budding yeast have an additional known MT depolymerase, Kip3, which is a kinesin-8 (Gupta et al., 2006). Based on our data presented here and on our previous work in which we quantitatively analyzed MT dynamics in the absence of one or both of these depolymerases, it seems possible that removing these proteins from MTs creates new opportunities for other MAPs to affect assembly and disassembly rates (Bergman et al., 2019). Bim1, the EB1 homolog, is responsible for promoting MT growth (Blake-Hodek et al., 2010; Wolyniak et al., 2006). When one of its binding partners (e.g. Kar3Cik1) is absent, there may be more protein available to increase growth rates. It may also be that these proteins act as inducers of catastrophe, but could possibly modulate the rate at which MTs depolymerize once a catastrophe has occurred.

Locomotion of Kar3 heterodimers in either direction on MTs in our *in vitro*, single-molecule assay was rare and was never observed in our time-lapse imaging of live cells. The plus-end directed motility we observed *in vitro* was variable in velocity and duration, and its mechanism is unclear. In contrast, the minus-end directed motility we observed during metaphase was more uniform in velocity, matching the speed of captured kinetochores being transported along kMTs in cells in an inducible chromosome capture system (Tanaka et al., 2005). Our calculated minus-end directed velocity of 0.64 ± 0.27 μm min^-1^ was ∼7-fold slower than velocities (77 nm s^-1^ (4.6 μm min^-1^) reported by others for Kar3Cik1 (Mieck et al., 2015). Mieck and colleagues reported that Kar3Vik1 moves at speeds approaching 234 nm s^-1^(∼14 μm min^-1^), a speed which we would not observe at our sampling rate. Such a high rate might explain our failure to observe Kar3 movement along cMTs in cells.

### Differential Cell Cycle Regulation of Kar3 Heterodimers

By using a combination of cell cycle arrests and acute, targeted depletion of accessory proteins, we show at a subcellular level where heterodimer populations function on MTs throughout the cell cycle. Fluorescently-tagged Kar3 fusion proteins have been used previously to ascertain where Kar3 heterodimers are located in cells (Gardner et al., 2008; Gibeaux et al., 2006; Maddox et al., 2003; Mieck et al., 2015), but have not focused on how localization transitions from one cell cycle stage to the next. Recent work showed that Cik1 or Vik1 absence changes Kar3-GFP localization on anaphase spindles (Mieck et al., 2015). We confirmed these previous observations and extended this analysis using a GFP-Kar3 construct in strains that could be arrested at different cell cycle stages (Fig. 1A). We then determined which populations were Kar3Cik1 vs Kar3Vik1 by tagging one of the accessory proteins with mScarlet-I (Fig. 1B and C).

These data, combined with analysis of strains expressing a degron-tagged version of an accessory protein and GFP-Kar3, independently confirmed that Kar3Cik1 resides on the nuclear face of SPBs, along the spindle, and on kMTs (Fig. 2A), and that Kar3Vik1 is on the cytoplasmic face of SPBs and on cMTs plus-ends (Fig. 2B). We also confirmed that Kar3’s interaction with MTs is dependent upon heterodimer formation (Fig. 2C) and that the accessory proteins cannot bind MTs in the absence of Kar3 (Fig. S1) (Manning et al., 1999).

Our *in vivo* localization data are consistent with our single-molecule observations. The most common Kar3-MT interaction we observed was plus-end tracking, which was highly, but not completely, dependent on Cik1 (Table 5). This observation aligns well with the low frequency of Kar3Vik1 we observed on cMT plus ends and the relatively high frequency of Kar3Cik1 on kMTs in cells (Fig. 1). Association of Kar3 with MT minus-ends in our assay was the second most common observation. This interaction was also dependent on heterodimer formation, but more so on Vik1 than Cik1 (Fig. 5). Consistently, SPB staining in cells was only slightly diminished in Cik1-depleted cells, but was nearly absent in the Vik1-depleted cells.

Previous analysis of *kar3Δ* phenotypes revealed an increase in cMT number and length (Huyett et al., 1998; Saunders et al., 1997b). We also observed an increase in cMT numbers for Kar3- and Vik1-depleted cells at all cell cycle stages (Fig. 3C). However, the average length of cMTs was only increased in G1 for Kar3-, Cik1-, and Vik1-depleted cells, and in anaphase-arrested cells depleted of Vik1 (Fig. 3D-F). Our results may differ from previous observations because of one or more of several differences in the set up. Instead of observing asynchronous cells carrying gene disruptions, we observed arrested cells depleted of a protein of interest by use of a degron tag. Perhaps the increase in cMT length observed in earlier studies was the result of the cumulative effect of the protein’s absence over multiple cell cycles in a null mutant. Mutations in *KAR3* and *CIK1* both lead to shorter metaphase spindles that collapse or break as mitosis progresses. Our data suggest that these phenotypes arise during metaphase and that Kar3Cik1 is necessary for building a spindle of appropriate length and for maintaining the necessary opposing force across the spindle (Figs. 2A, C, 3A and S1C). Long spindles can eventually form in *cik1Δ* cells (Gardner et al., 2008; Mieck et al., 2015). According to our data, Kar3Cik1 is necessary for maintaining proper length of spindle during anaphase (Figs. 2A, 3B). Vik1 had not previously been implicated in spindle length maintenance, but we observed a statistically significant increase in the spindle length during late anaphase in Vik1-depleted cells (Fig. 3B). It is possible that the increased number of cMTs in these cells (Fig. 3C) may exert a greater force on SPBs to cause this. This result may reflect a previously unknown function of Kar3Vik1.

### Toward an Integrated Understanding of Kar3 Function and Regulation

Though much work has been done investigating Kar3 activity in the cell, most of this work focused on the Kar3Cik1 heterodimer. Here we have dissected apart the distinct localizations and functions of the two different Kar3 heterodimers, and have uncovered previously unknown functions of Kar3Vik1. Our reconstituted MT dynamics assay implicated Kar3Vik1 as a modulator of MT dynamics and a promoter of catastrophes. Localization data from our studies and those of others revealed that Kar3Vik1 is present exclusively in the cytoplasm, which implies that these activities affect only cMTs. Overall, Kar3Vik1 may primarily act to limit cMT length and numbers, particularly during anaphase (but perhaps all of mitosis) to help control spindle length and to help efficiently orient the nucleus for efficient chromosome inheritance.

Across species, kinesin-14s serve a large variety of functions. The largest number of the kinesin-14s identified to date are found in land plants, which lack cytoplasmic dynein (Gicking et al., 2018). These land plant kinesin-14s presumably had to expand to fulfill all the roles normally performed by dynein. Other examples of kinesin-14s include Drosophila Ncd (McDonald et al., 1990), mammalian HSET (Mountain et al., 1999), and fission yeast pkl1 and klp2 (Pidoux et al., 1996; Troxell et al., 2001). These kinesin-14s differ from budding yeast Kar3 because they act as homodimers. Their functions seem to be balancing forces within the spindle and capturing kinetochores, similar to Kar3Cik1. The complexity of Kar3 activities described here through the cell cycle, suggest that budding yeast might use their kinesin-14, Kar3, for many of the same functions as in these other organisms. However, due to a closed mitosis, yeast have separate nuclear and cytoplasmic microtubule cytoskeletons. They also are unique in use of different kinesin-14 accessory proteins, Cik1 and Vik1, in the nucleus and the cytoplasm, respectively. Having separate Kar3populations with different accessory factors allows budding yeast to compartmentalize microtubule functions and cell cycle regulation.

## MATERIALS AND METHODS

### Yeast Strains, Culturing and Harvesting

All *Saccharomyces cerevisiae* strains in this study are s288c background and can be found in Table S1. Fluorescent and degradation tags were integrated into the genome by homologous recombination as previously described. Strains were grown in standard rich medium (YPD) at 25°C. Strains containing AID tag alleles were treated with final concentrations of 250 µM 3-indole acetic acid (Sigma-Aldrich, St. Louis, MO, USA) in DMSO and buffered with 50 mM potassium phosphate buffer at pH 6.2 for the 30 minutes prior to harvesting or imaging. Lysates were prepared with a SPEX cryogenic grinder (SPEX Sample Prep, Metuchen, NJ, USA) as described in (Bergman et al., 2019).

### Live-Cell Imaging of Yeast

Strains were grown overnight in rich media at permissive temperature. They were then diluted to OD = 0.1 in casamino acid media (0.67% yeast nitrogen base with ammonium sulfate, 2% casamino acid, 2% glucose) supplemented with amino acids and grown at permissive temperature to recover. Cells were then shifted to 37°C for 3.5 hours. If necessary, 3-IAA was added in the last 30 minutes of arrest. Cells were then mounted onto coverslips coated with concanavalin A (Sigma-Aldrich, St. Louis, MO, USA) and covered with pre-warmed media. Microscopy was performed on a Nikon Eclipse Ti2 with a Nikon 60X Plan Apochromat objective and a 1.5X booster (Nikon, Tokyo, Japan) using an Orca Flash 4.0 camera (Hamamatsu, Shizuoka, Japan). Images were captured using Nikon Elements software. Temperature was maintained with a heated chamber and objective collar (Oko Labs, Pozzuoli, Italy). GFP was excited using a 488-laser line. mScarlet-I and mRUBY2 were excited using a 561-laser line. Eleven slice z-stack series spaced 0.5 µm apart were taken every 30 seconds for 10 minutes.

### Immunoblotting

To compare relative protein concentrations in lysates, equal amounts of thawed and diluted lysate powder were run on an 8% polyacrylamide gel. Preparation of cleared lysate is described in Bergman and Wong 2018. Total lysate was prepared by adding 220 µL of cold 2X PEM and 0.5 µL of Protease Inhibitor Cocktail IV (Roche, Basel, Switzerland) to 0.22 g of lysate powder. The mixture was solubilized with an equal volume of 2X SDS sample buffer, boiled for 1 minute, and briefly centrifuged before the supernatant was loaded onto a gel. Immunoblotting was performed with 1:750 9E10 mouse anti-myc primary antibody, 1:20,000 anti-PGK antibody, and 1:1000 rat anti-α-tubulin (Santa Cruz Biotechnology, Cat.# 5C-53030, Dallas, TX, USA). Band intensities were measured using Image Studio Lite (LI-COR, Lincoln, NE, USA).

### Glass slide cleaning and Cover Glass Passivation

Glass slides were washed with acetone and then 100% ethanol and left to air dry. Cover glasses were passivated by sonication in acetone for 30 minutes, followed by a 15 minute soak in 96% ethanol. The cover glasses were then rinsed three times with ddH_2_O and left in a 2% Hellmanex III solution for 2 hours. Afterwards, the coverglasses were rinsed three times with ddH_2_O and dried with nitrogen gas. Cover glasses were then laid over a 50 µL drop of 100 mM HEPES, 1 mg mL^-1^ PLL-g-PEG, and 1 mg mL^-1^ PLL-PEG-Biotin for 1 hour in a humidity chamber. Cover glasses were then rinsed with 1X PBS for 2 minutes and then in ddH_2_O for 2 min. Finally, they were dried with nitrogen and stored at 4°C.

### Preparation for TIRF Microscopy

Assembly of flow chambers for TIRF imaging was performed as in Bergman and Wong 2019. Similarly, generation of microtubule seeds followed (Bergman et al., 2019), with the exception of using HiLyte 647-labeled tubulin (Cytoskeleton Inc., Denver, CO, USA) instead of rhodamine-labeled tubulin.

### Preparation of Whole Cell Lysates

For MT dynamics assays, lysates were prepared as in (Bergman et al., 2019) except both exogenous ATP and GTP were excluded from the final clarified lysate. Instead, 30 µL of clarified lysate was directly loaded into the flow chamber.

For characterization of Kar3 on microtubules *in vitro*, lysates were prepared as above with the following modifications. To enrich the population of soluble Kar3, frozen lysate powder was not diluted with 10X PEM buffer. Instead, 0.5 µL of Protease Inhibitor Cocktail IV (Roche, Basel, Switzerland) was added to 0.24 g of lysate powder. The mixture was thawed on ice for 10 minutes and then sonicated for 2 seconds (1 second cycle time, 20% duty cycle, output control set to 2) using a W-385 sonicator with a #419 microtip (Heat Systems - Ultrasonics, Inc., Farmingdale, NY, USA) prior to ultracentrifugation.

### TIRF Microscopy

MT dynamics reconstitution assays were performed on a Nikon Eclipse Ti2 inverted TIRF microscope a Nikon 60X Plan Apochromat TIRF objective (NA 1.49) with a 1.5X booster lens (Nikon, Tokyo, Japan) in a temperature-controlled chamber (Oko Labs, Pozzuoli, Italy) warmed to 28°C and captured with a Orca Flash 4.0 camera (Hamamatsu, Shizuoka, Japan) operated by Nikon Elements software. Single-plane TIRF images of 488 nm and 561 nm illumination were taken every 5 seconds for 30 minutes.

Single-molecule studies of motors on microtubules were performed on Nikon Eclipse Ti2 inverted microscope using a Nikon 60X CFI Apo TIRF objective (NA 1.49) and an Orca Fusion Gen III sCMOS camera (Hamamatsu, Shizuoka, Japan) at 1.5X magnification using the Nikon NIS Elements software. Using a LUNF 4-line laser launch (Nikon Instruments, Tokyo, Japan) and an iLas2 TIRF/FRAP module (Gataca Systems, Massy, France) total internal reflection fluorescence (TIRF)

### Image and Data Analysis

Image files were analyzed using Fiji (NIH). For finding an average spindle length, 50 spindles from either metaphase- or anaphase-arrested genotypes were counted. The average lengths of cytoplasmic MTs and the average number of cytoplasmic MTs per cell was calculated by counting and measuring the length and number of visible MTs not in the nucleus from 50 cells at each condition at the zero time-point. Statistical significance was determined by one-way ANOVA using the Kruskal-Wallis test comparing the depleted strains to the WT. Kymographs for measuring dynamics were constructed from all MTs whose entire length was trackable for the entire 30-minute observation window after registration. Dynamics parameters and statistics were calculated as in (Bergman et al., 2019).

## ACKNOWLEDGMENTS

The authors wish to thank Julia Torvi for very helpful discussion, construction of the GFP-KAR3 allele and strain, and aid in designing experiments. We also thank members of the Barnes/Drubin lab for discussion of experiments and analysis. The mRUBY2-TUB1 constructs were a gift from W. Lee (Dartmouth University).

---

**Figure S1.**
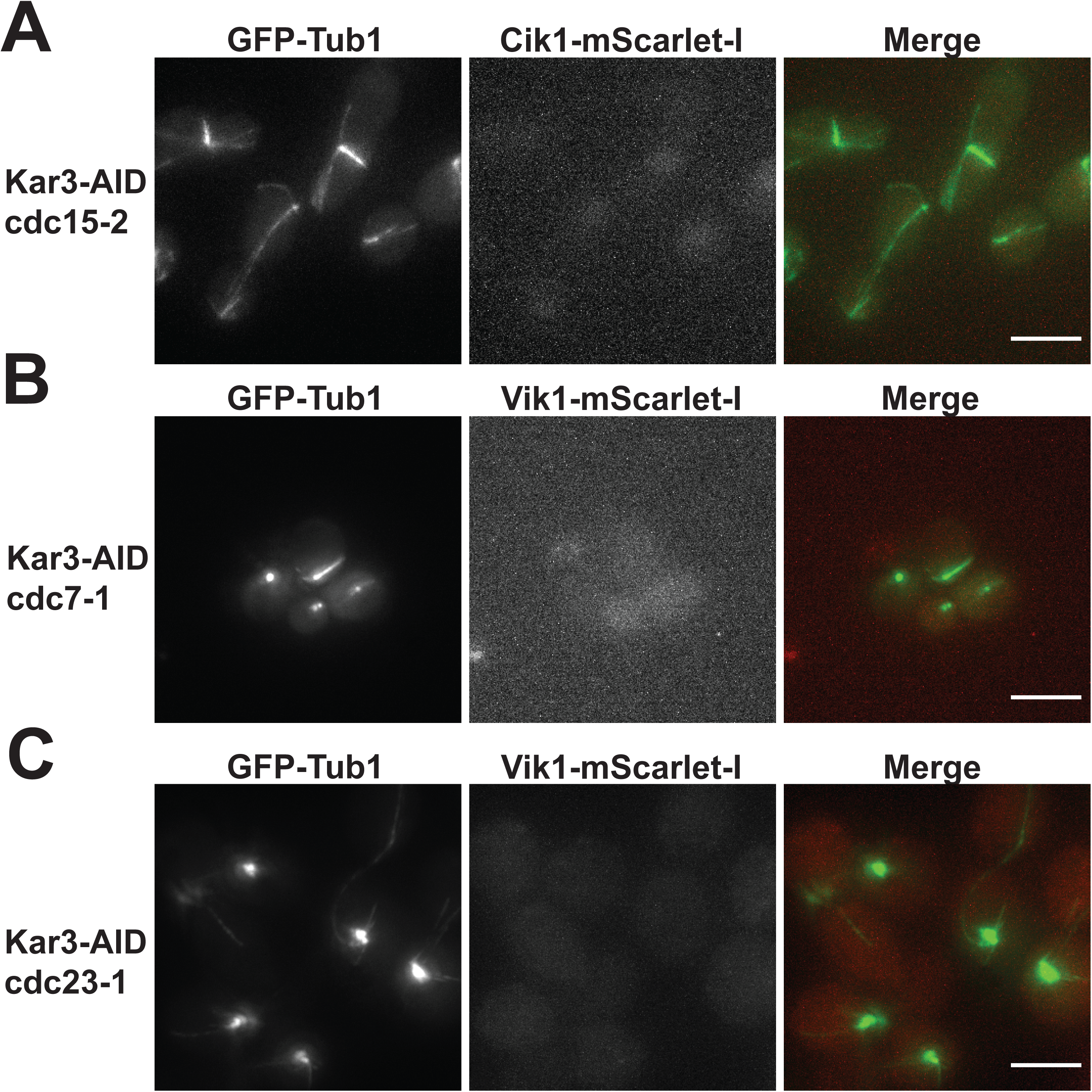
Accessory Protein Localization Is Dependent Upon Kar3. GFP-TUB1 KAR3-AID cells were arrested and depleted of Kar3. **(A)** Cik1-mScarlet-I is diffuse in anaphase-arrested cells depleted of Kar3. **(B)** Vik1-mScarlet-I localization in S phase without Kar3. **(C)** Vik1-mScarlet-I localization and GFP-tubulin morphology in Kar3-depleted cells arrested in metaphase. Scale bars are 5 µm.

**Figure S2.**
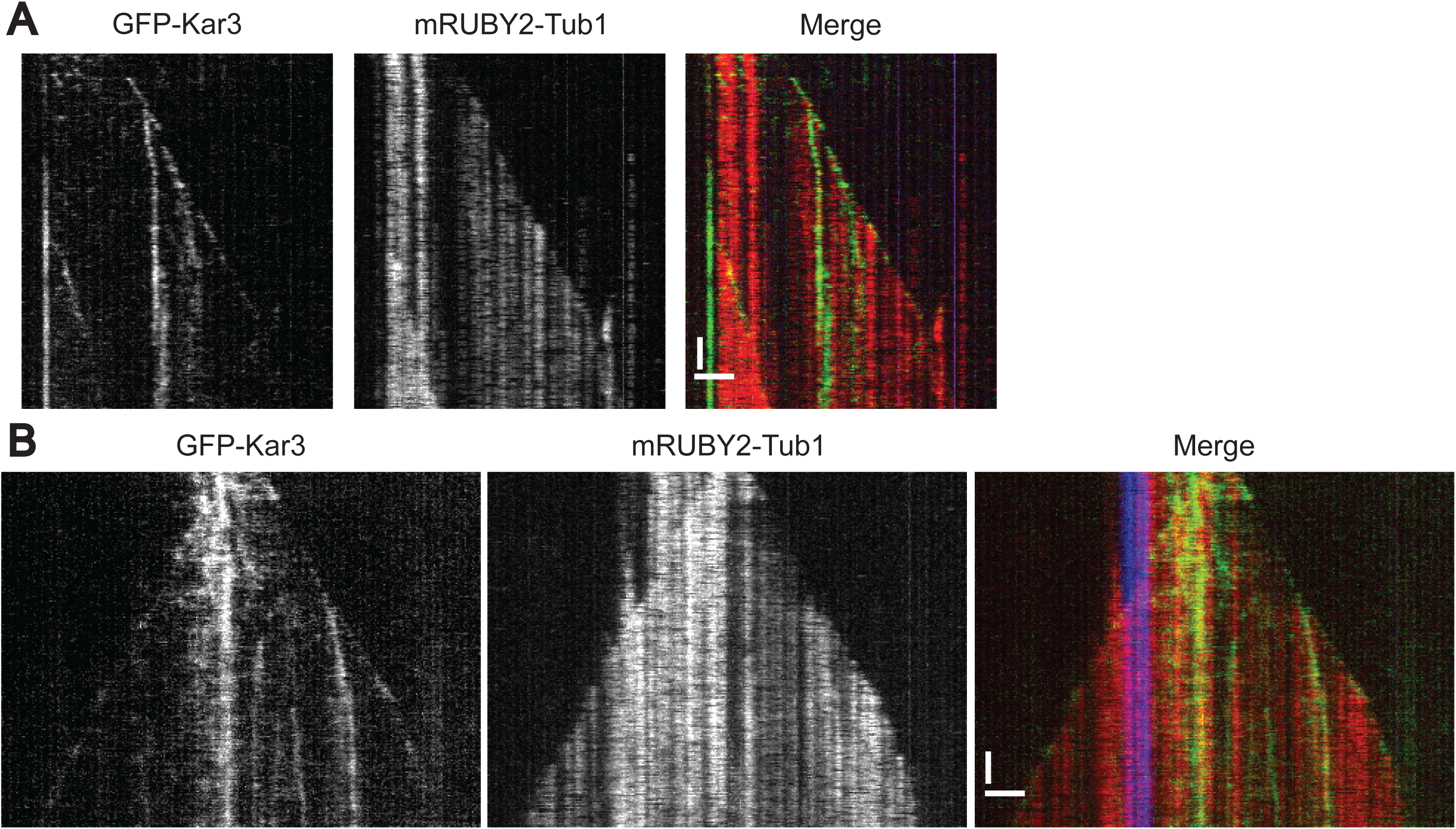
Non-Motile Kar3 Diffusely Covers Coats MTs in the Absence of Cik1 During Anaphase. Dynamic MTs were reconstituted from lysate arrested in anaphase and depleted of Cik1. GFP-Kar3 (green), mRUBY2-TUB1 (red), and HiLyte647-labeled seeds (blue). **(A)** GFP-Kar3 associated with a single MT. **(B)** GFP-Kar3 associated with an anti-parallel MT. Scale bars are 2 µm and vertical bars are 2 min.

**Table S1.**
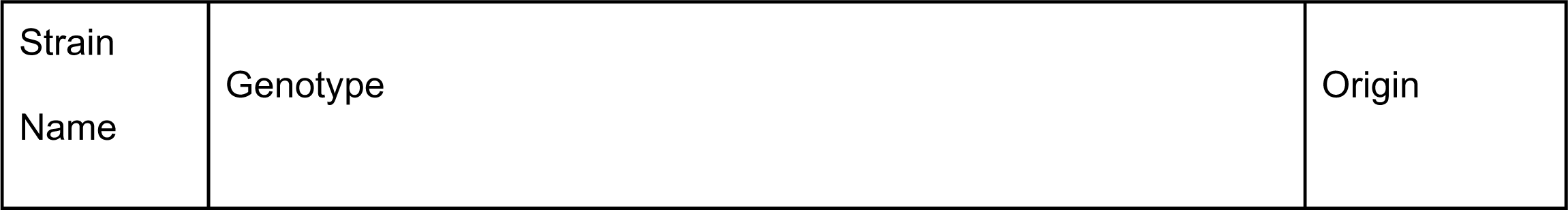
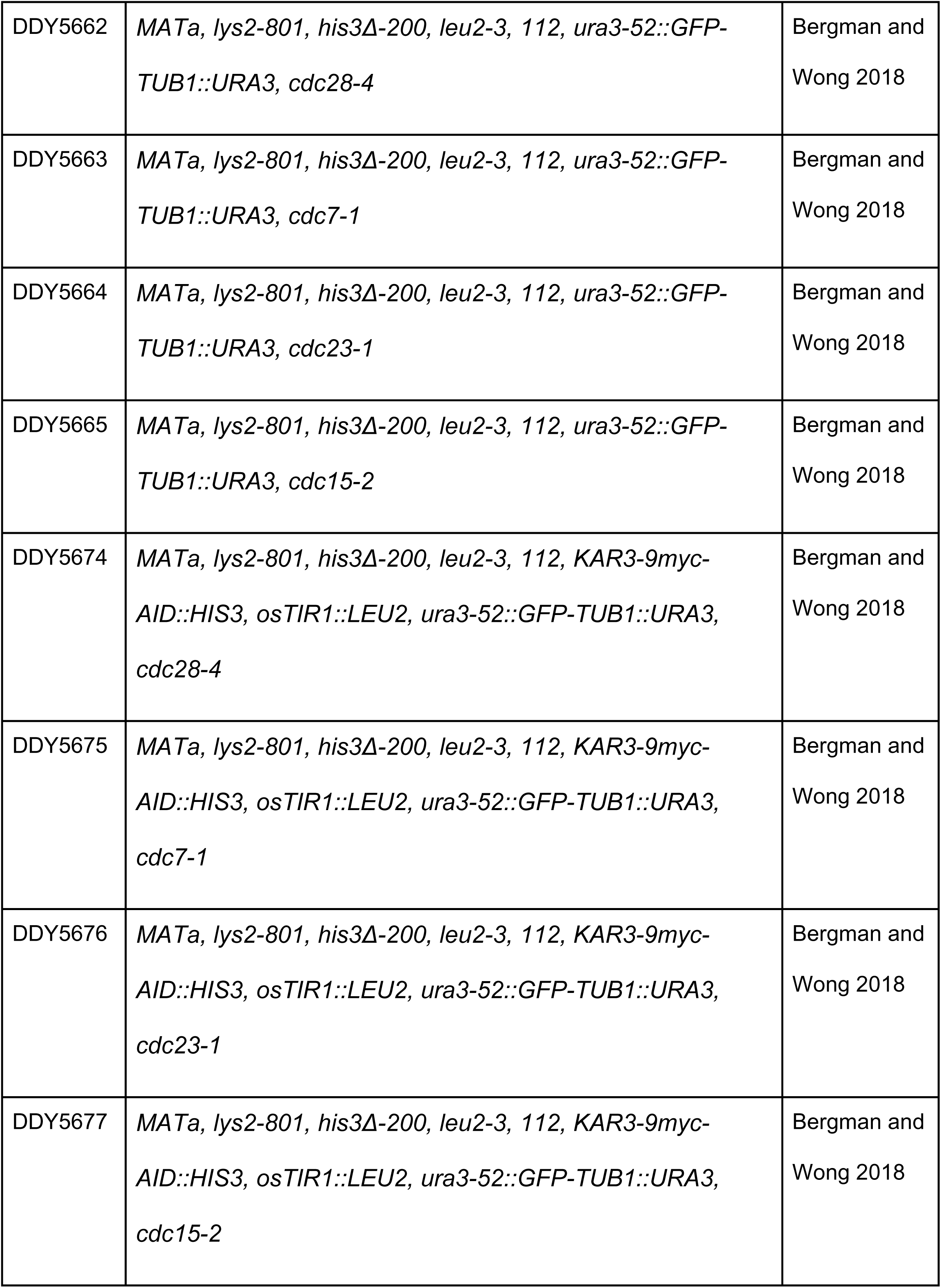
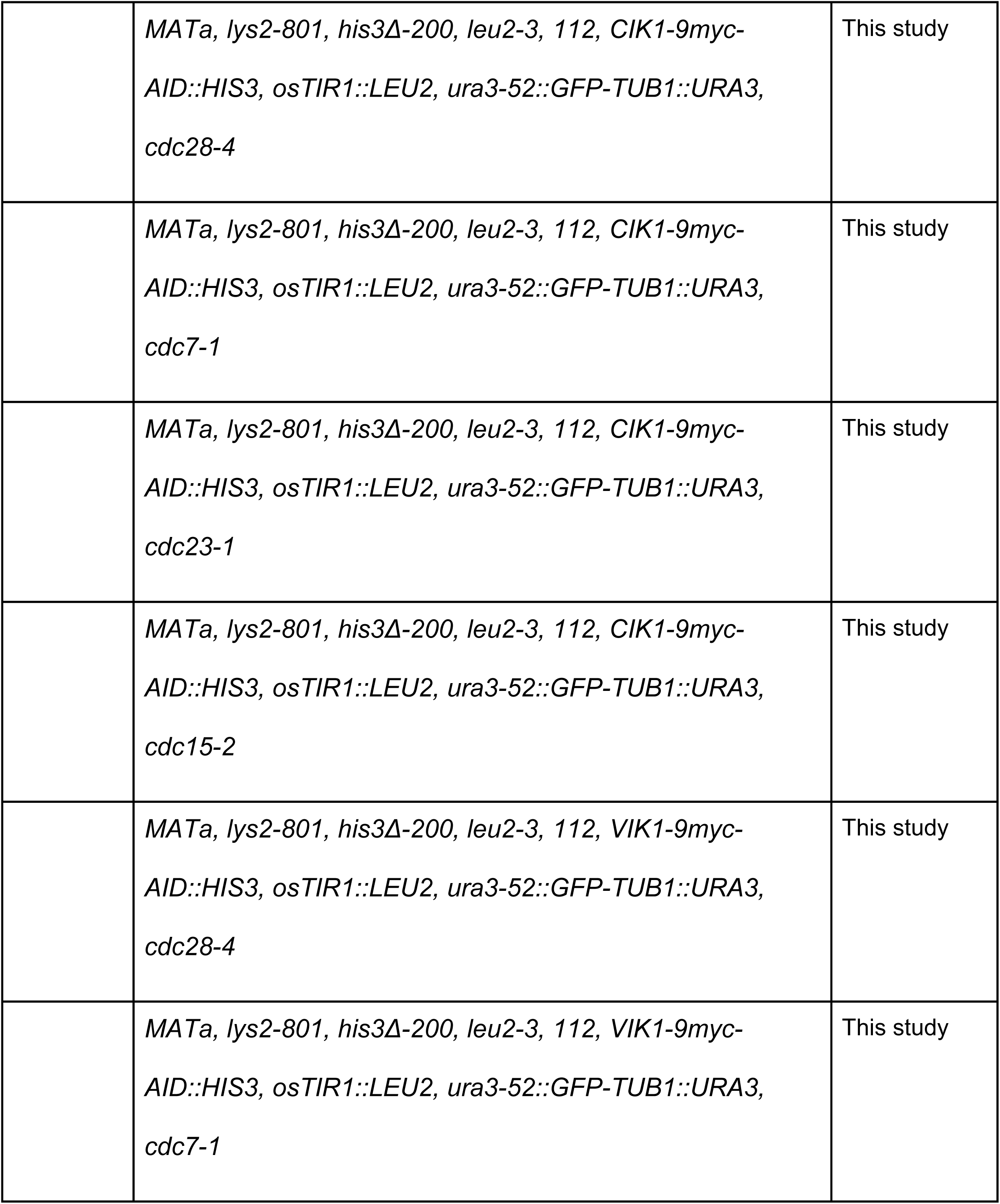
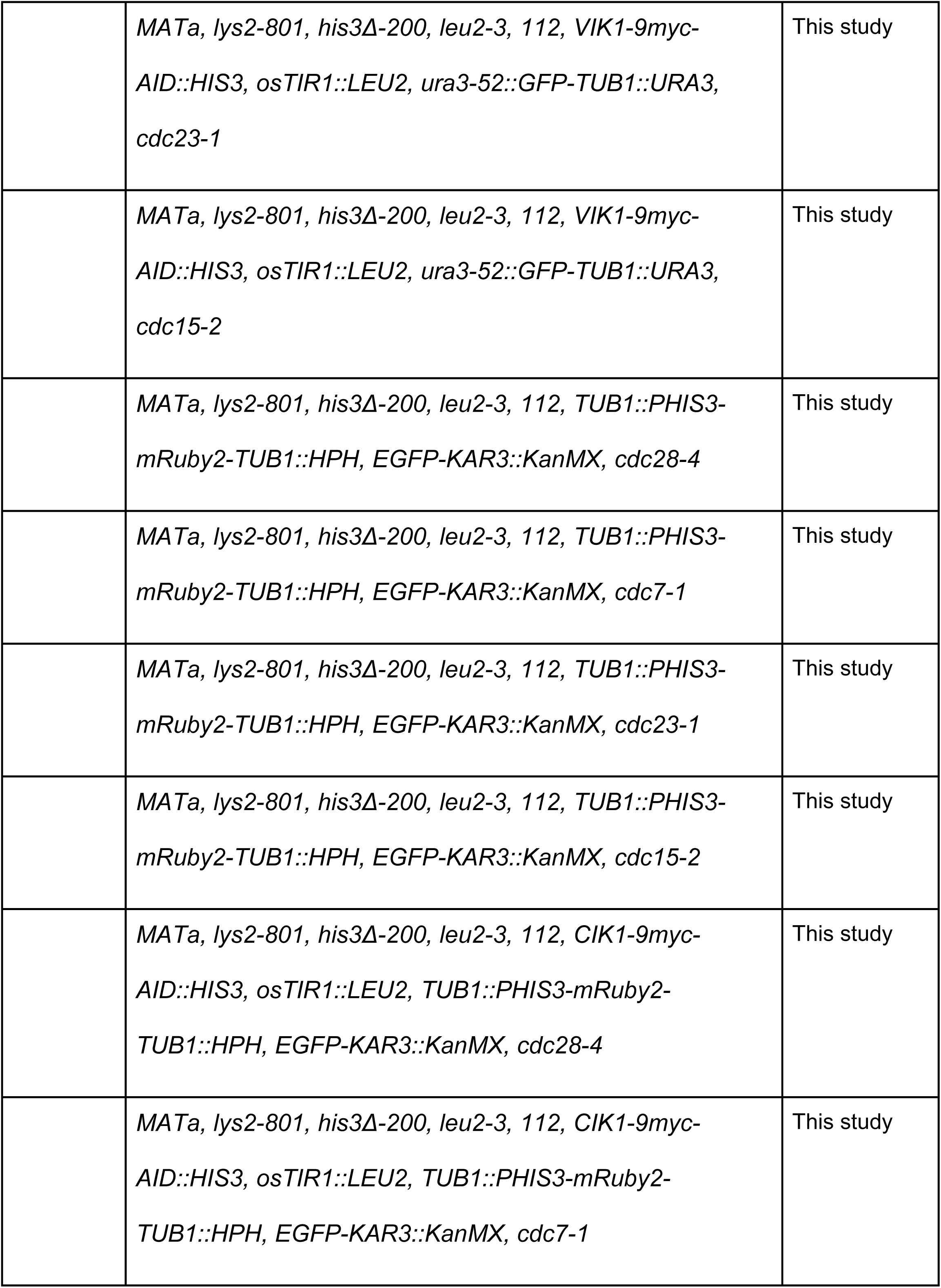
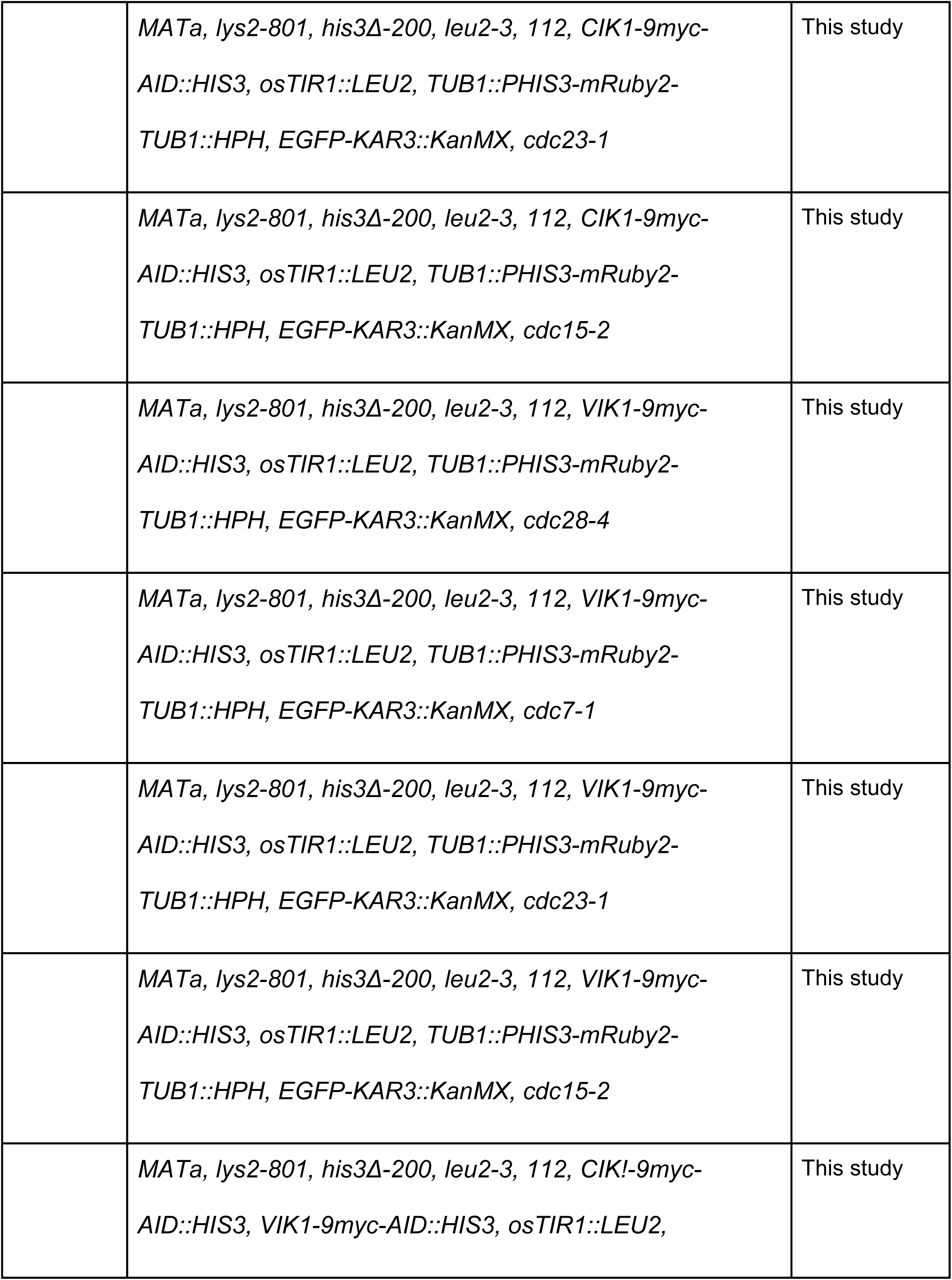
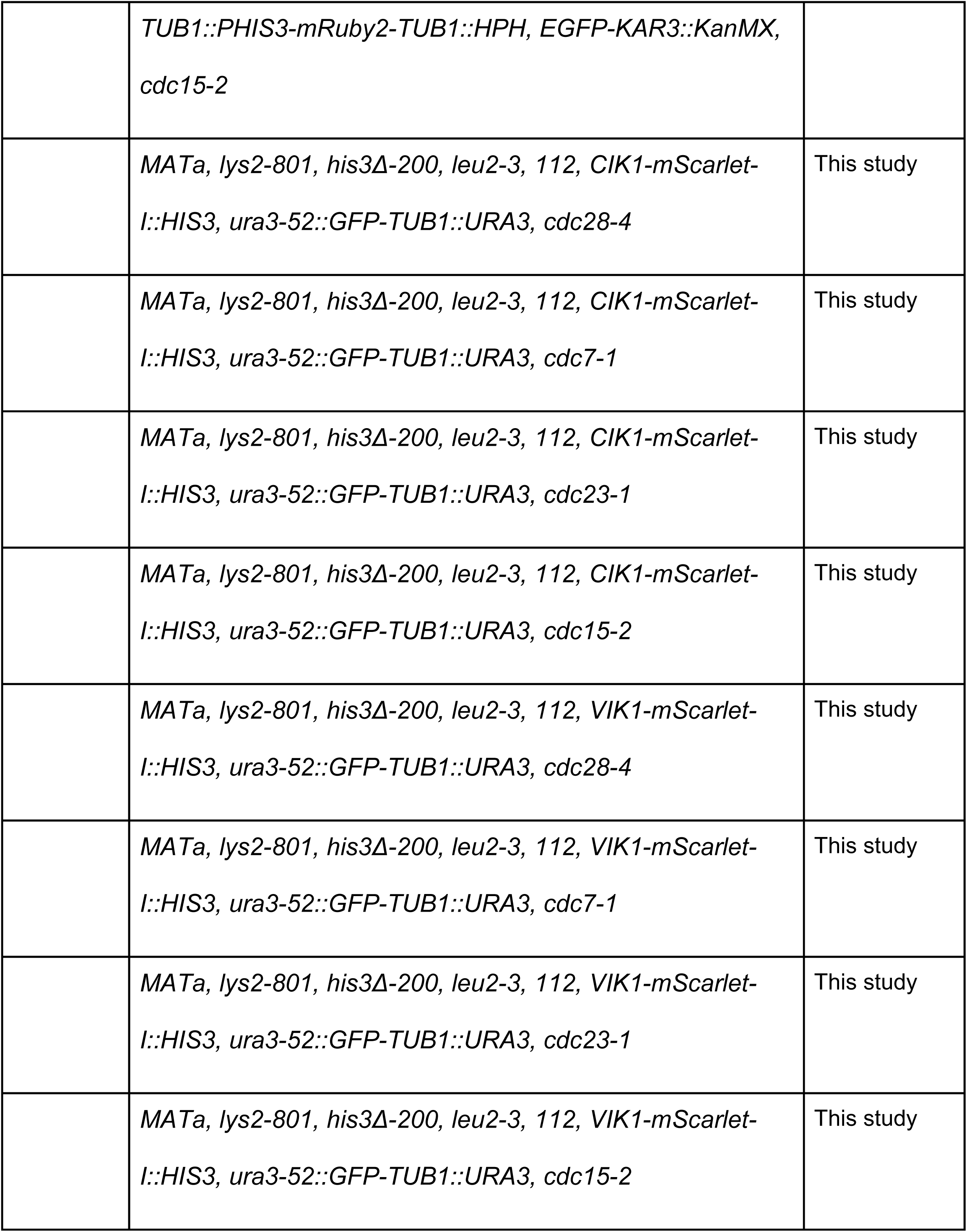
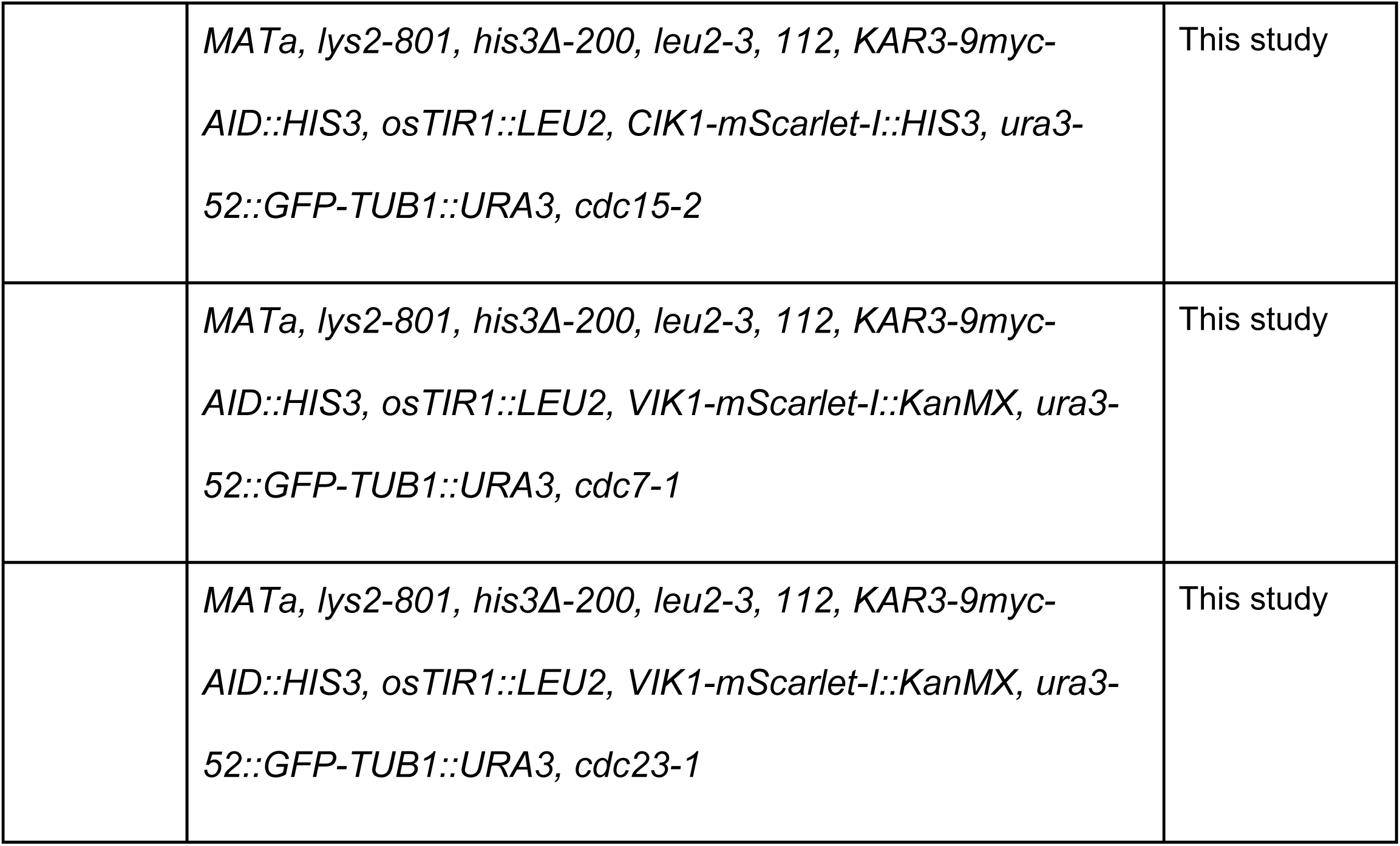
Strains Used in this Study.

